# Heme allocation in eukaryotic cells relies on mitochondrial heme export through FLVCR1b to cytosolic GAPDH

**DOI:** 10.1101/2024.04.23.590782

**Authors:** Dhanya Thamaraparambil Jayaram, Pranav Sivaram, Pranjal Biswas, Yue Dai, Elizabeth A. Sweeny, Dennis J. Stuehr

## Abstract

Heme is an iron-containing cofactor essential for life. In eukaryotes heme is generated in the mitochondria and must leave this organelle to reach protein targets in other cell compartments. Mitochondrial heme binding by cytosolic GAPDH was recently found essential for heme distribution in eukaryotic cells. Here, we sought to uncover how mitochondrial heme reaches GAPDH. Experiments involving a human cell line and a novel GAPDH reporter construct whose heme binding in live cells can be followed by fluorescence revealed that the mitochondrial transmembrane protein FLVCR1b exclusively transfers mitochondrial heme to GAPDH through a direct protein-protein interaction that rises and falls as heme transfers. In the absence of FLVCR1b, neither GAPDH nor downstream hemeproteins received any mitochondrial heme. Cell expression of TANGO2 was also required, and we found it interacts with FLVCR1b to likely support its heme exporting function. Finally, we show that purified GAPDH interacts with FLVCR1b in isolated mitochondria and triggers heme transfer to GAPDH and its downstream delivery to two client proteins. Identifying FLVCR1b as the sole heme source for GAPDH completes the path by which heme is exported from mitochondria, transported, and delivered into protein targets within eukaryotic cells.

## Introduction

Life evolved to widely utilize a special form of iron that is bound within the protoporphyrin IX ring (iron protoporphyrin IX; heme) ^1,2^. Heme biosynthesis in eukaryotes is a complex process and its final steps occur inside the mitochondria ^3^. However, how heme is then exported from mitochondria, transported, and inserted into numerous proteins that require heme for function but reside elsewhere in cells has been unclear. Because heme is chemically reactive and has promiscuous binding properties, its synthesis is tightly controlled and its intracellular transport has long been imagined to involve macromolecular carriers ^2,4,5^. Of all the proteins or other macromolecules that have been proposed, the protein glyceraldehyde phosphate dehydrogenase (GAPDH), an enzyme in the glycolytic pathway that is ubiquitously expressed and known to perform alternative moonlighting functions ^6–8^, has recently emerged as a premier intracellular heme chaperone ^9^, based on findings that GAPDH binding of mitochondrially-generated heme is required for and coupled to intracellular heme delivery to numerous targets including hemoglobins α, β, and γ ^10^, myoglobin ^10^, nitric oxide synthases ^11–13^ soluble guanylyl cyclase β-subunit (sGCβ)^14^, cytochromes P450^15^, heme oxygenase 2 ^16^, indoleamine dioxygenase 1 (IDO1) and tryptophan dioxygenase (TDO) ^17^. Insertion of the GAPDH-sourced heme into recipient target proteins is the final downstream step in heme delivery, and is now understood to require the cell chaperone protein Hsp90, which is typically bound to the heme-free (apo-) forms of the recipient proteins and drives their heme insertions in an ATP-driven process ^18^. Thus, GAPDH binding of mitochondrial heme enables its wide distribution in cells, most often capped by an Hsp90-assisted heme insertion into the protein target. However, one key question remains unanswered in the overall pathway, namely, how does mitochondrial heme reach GAPDH? Here, we address this question in experiments that utilize a cultured human cell line, isolated mouse mitochondria, and a newly described GAPDH reporter protein ^19^ whose heme binding in living cells can be followed in real-time based on the change in its fluorescence intensity. Our results identify a transporter protein located in the outer mitochondrial membrane, Feline Leukemia Virus subgroup C Receptor 1b (FLVCR1b), as the sole conduit for mitochondrial heme export to GAPDH in the cells, operating through a mechanism that involves a direct FLVCR1b-GAPDH interaction for heme exchange. Overall, our findings elucidate the route of heme export from the mitochondria and show how this action is tightly coupled to heme distribution and insertion into final destination proteins inside eukaryotic cells.

## Results

### FLVCR1b supplies heme to GAPDH and enables downstream heme delivery

To explore how heme that is generated in mitochondria transfers to GAPDH in the cell, we tested the importance of FLVCR1b, a transmembrane protein exclusively localized in the outer mitochondrial membrane that has previously been implicated in mitochondrial heme export in erythrocytes during erythropoiesis ^20^. We performed experiments in a human embryonic kidney cell line (HEK293T cells) transfected to express an HA-tagged human GAPDH reporter protein called TC-hGAPDH ^19^, which after being labeled with FlAsH reagent ^19,21^ signals its heme binding in living cells or in solution by a fluorescence quenching of its FlAsH signal ^19^. We determined how siRNA-targeted knockdown of cell FLVCR1b expression impacts heme loading onto TC-GAPDH in living cells in response to our stimulating their mitochondrial heme biosynthesis by providing the heme precursor molecules δ-aminolevulinic acid (δ-ALA) and ferric citrate (Fe) ^22^, which we have previously shown causes the total heme level in the HEK293T cells, and the level of heme bound in the TC-hGAPDH, to increase by about 3-fold within 2 h ^19^. The targeted siRNA treatment diminished cell FLVCR1b protein expression relative to a scrambled siRNA control by 70 +/- 9% (n = 5) (Fig. 1A and S1A) and this did not impact the cell heme level from increasing in response to the δ-ALA/Fe addition (Table S1), consistent with a previous report ^20^. However, the FLVCR1b knockdown inhibited transfer of mitochondrial heme into TC-hGAPDH by more than 90%, as indicated by the TC-hGAPDH fluorescence signal in the FLVCR1b knockdown cells remaining stable with time after the δ-ALA/Fe addition, compared to the large decrease in TC-hGAPDH fluorescence that occurred in similarly-treated cells given the scrambled siRNA (Fig. 1B). The FLVCR1b knockdown did not alter cell expression of TC-hGAPDH (Fig. S1B) nor did it inhibit TC-hGAPDH from obtaining heme when it was provided externally to the cell cultures (Fig. S2A and B). Thus, a targeted knockdown of FLVCR1b expression greatly restricted mitochondrial heme transfer to TC-hGAPDH in the living cells.

**Fig. 1:**
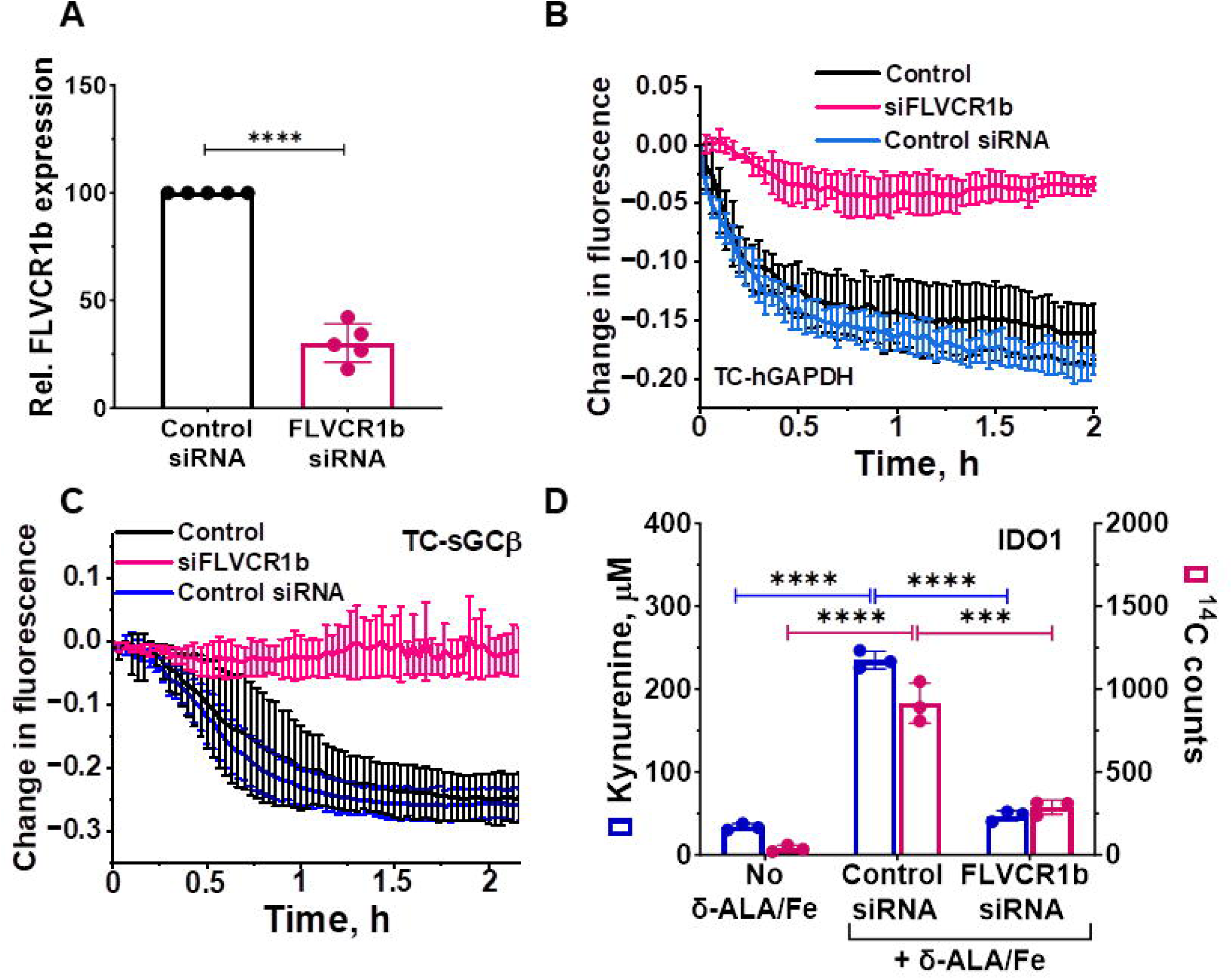
FLVCR1b mediates mitochondrial heme transfer to GAPDH and enables its downstream heme delivery in cells. **A** Targeted siRNA-based knockdown of cell FLVCR1b expression in HEK293T cells. Mean +/-SD of 5 independent trials. **B** Knockdown of cell FLVCR1b expression inhibits mitochondrial heme transfer to FlAsH-labeled TC-hGAPDH in living cells whose mitochondrial heme biosynthesis was stimulated by δ-ALA/Fe addition at time = 0. Representative of three independent trials, mean +/- SD of triplicates. **C**, **D** FLVCR1b knockdown inhibits the delivery of mitochondrial heme to two GAPDH client proteins in the living cells after the δ-ALA/Fe addition, as judged by **C** it blocking the decrease in fluorescence intensity for FlAsH-labeled TC-sGCβ and **D** it blocking the increase in the enzymatic activity and ^14^C heme content of IDO1. **C**, Representative of three independent trials, mean +/- SD of triplicates. **D**, Mean +/- SD of 3 independent trials. Significance: *** p < 0.001 and **** p < 0.0001 vs. the compared group based on a one-way ANOVA test.

We next examined how the FLVCR1b knockdown would impact downstream delivery of mitochondrial heme to two protein targets that we expressed separately in the cells and that are known to have GAPDH-dependent heme deliveries (a TC-tagged version of soluble guanylyl cyclase β-subunit termed TC-sGCβ, and indoleamine dioxygenase-1, IDO1) ^14,17^. The FLVCR1b knockdown did not alter cell expression of the TC-sGCβ and IDO1 target proteins (Fig. S1C and D) but it did completely block their heme deliveries, which otherwise took place following δ-ALA/Fe addition (Fig. 1C and D). Thus, the mitochondrial heme exporter FLVCR1b is needed for GAPDH to obtain mitochondrial heme in the cells, and this in turn is required for delivery of the mitochondrial heme to two GAPDH-dependent protein targets, revealing that in the absence of a FLVCR1b-GAPDH heme transfer the cells had no alternative way to accomplish the heme deliveries.

### FLVCR1b and GAPDH directly interact during mitochondrial heme export

We next investigated if mitochondrial heme export to GAPDH involved its interaction with FLVCR1b, whose expression is restricted to the outer mitochondrial membrane ^20,23^. We utilized the Duolink proximity ligation assay (PLA) which detects the interaction of two proteins within a 0-40 nm distance, a typical range for protein partners that engage in direct interaction ^24^. We compared the extent of FLVCR1b interaction with GAPDH before and after providing cells with δ-ALA/Fe to stimulate their mitochondrial heme biosynthesis and to enable the consequent heme loading into GAPDH as shown above. Using the kinetic data from Fig. 1B as a guide, we assessed the level of GAPDH-FLVCR1b interaction at 0, 15, 30, 45, 60, and 120 min after adding δ-ALA/Fe to the cells, which correspond to times before, during, and after mitochondrial heme loading into GAPDH takes place. The PLA data in Fig. 2A and B show there was a detectable level of GAPDH and FLVCR1b interaction present in the resting cells (2.5x greater than background signal). Following the δ-ALA/Fe addition their interaction increased within 15 min and by 30 min had increased by 18-fold, and then gradually fell back to the original level by 60 min. Antibody pulldown experiments were done to independently assess the PLA results and they confirmed there is a temporal buildup in GAPDH-FLVCR1b association that peaked at 30 min after the δ-ALA/Fe addition (Fig S3A and B). In separate PLA-based experiments, we found that knockdown of FLVCR1b expression in the cells prevented GAPDH interaction with the mitochondrial surface protein HADHA and the increase in their interaction that otherwise occurred in response to δ-ALA/Fe addition in the control cells (Fig. 2C and D). This confirmed that the changes in GAPDH interaction specifically involved its interaction with mitochondrial FLVCR1b.

**Fig. 2:**
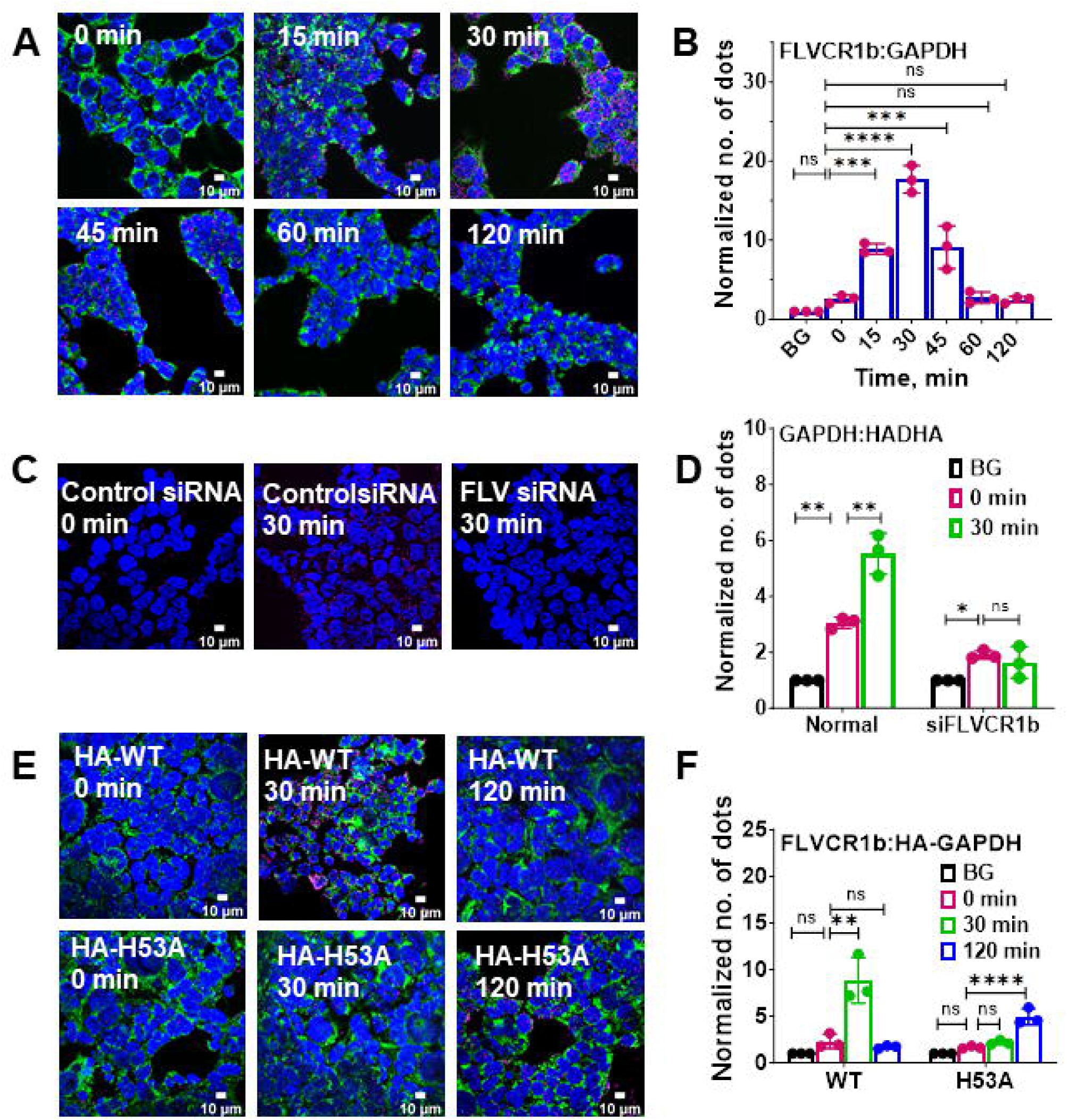
FLVCR1b interacts directly with GAPDH in a temporal manner during mitochondrial heme transfer in cells. Proximity ligation assay (PLA) was used to assess GAPDH interaction with FLVCR1b or with mitochondrial surface protein HADHA over time after addition of δ-ALA/Fe to cells to stimulate mitochondrial heme biosynthesis. **A** Representative fluorescence microscope images of cells stained with DAPI for nuclei (blue), using HADHA antibody for mitochondria (green), and to show the GAPDH-FLVCR1b interaction by PLA (red). **B** Quantification of the PLA results shows a temporal increase and decrease in the GAPDH-FLVCR1b interaction taking place following δ-ALA/Fe addition. BG, the background PLA signal level. **C** As in A, but with no mitochondrial staining and the FLVCR1b interaction with mitochondrial protein HADHA shown by PLA. **D** Quantification of PLA results showing that FLVCR1b knockdown abolished the increase in the GAPDH interaction with HADHA. **E** as in A, but with the PLA indicating FLVCR1b interaction with HA-tagged versions of GAPDH wild type or the heme binding-defective variant H53A GAPDH. **F** Quantification of PLA results showing that the temporal increase in FLVCR1b-HA-GAPDH interaction was muted and delayed for the H53A variant that cannot bind heme. **B, D, F** Mean +/- SD of 3 independent trials. Significance: * p < 0.05, ** p < 0.01, *** p < 0.001, and **** p < 0.0001 vs. the compared group based on a one-way ANOVA test. ns, not significant.

We further investigated the basis for the rise and fall in FLVCR1b-GAPDH interaction by expressing the H53A point mutant variant of HA-hGAPDH, which due to it having a 30-fold poorer heme binding affinity, ^12^ does not accumulate mitochondrially-generated heme in cells that are given δ-ALA/Fe ^12,19^. The PLA data in Fig. 2E and F show that there was a detectable level of H53A HA-GAPDH interaction with FLVCR1b present in the resting cells (about 2x above background), and upon δ-ALA/Fe addition we observed a delayed and partly muted rise in their interaction relative to what took place in replica experiments using cells expressing wild-type HA-GAPDH (Fig. 2E and F). Immunoprecipitation experiments confirmed the PLA-based results (Fig S4A-C). This suggests that heme binding to GAPDH was coupled to the rise and fall in its interaction with FLVCR1b that occurred when cell mitochondrial heme biosynthesis is stimulated.

### FLVCR1b in isolated mitochondria interacts with GAPDH to mediate heme export and delivery to target proteins

We next performed experiments with freshly isolated mitochondria from mouse brain and liver to expand on our cell culture findings. Upon incubating the isolated mitochondria for 30 min with mitochondria-free cell supernatant and δ-ALA and then reisolating the mitochondria, we observed a 3-fold increase in their heme level (Fig. 3A & B), confirming earlier reports that *in vitro* heme biosynthesis takes place in this circumstance ^25^. PLA experiments done with the re-isolated mitochondria showed that those that underwent the 30 min incubation with δ-ALA and cytosol had a four-fold increase in FLVCR1b-GAPDH interaction (Fig. 3C & D), mirroring what took place in living cells upon δ-ALA/Fe addition. In follow-up experiments we incubated the isolated mitochondria for 30 min with cytosol and ^14^C-δ-ALA so they would generate and accumulate radiolabeled ^14^C-heme, and then re-isolated these mitochondria for experiments. Upon placement in buffer solution at 37 C, the mitochondria released an inconsequential amount of ^14^C-heme into the solution over a 60 min observation period, as indicated by the ^14^C counts in the solution and mitochondria both remaining steady with time (Fig. 3E & F). However, the addition of purified GAPDH to these mitochondria resulted in an immediate and time-dependent export of the ^14^C-heme out of the mitochondria and into the solution, whereas adding a comparable molar amount of glutathione S-transferase, which had previously been reported to stimulate mitochondrial heme release ^26,27^ caused much less ^14^C-heme release (Fig. 3E & F).

**Fig. 3:**
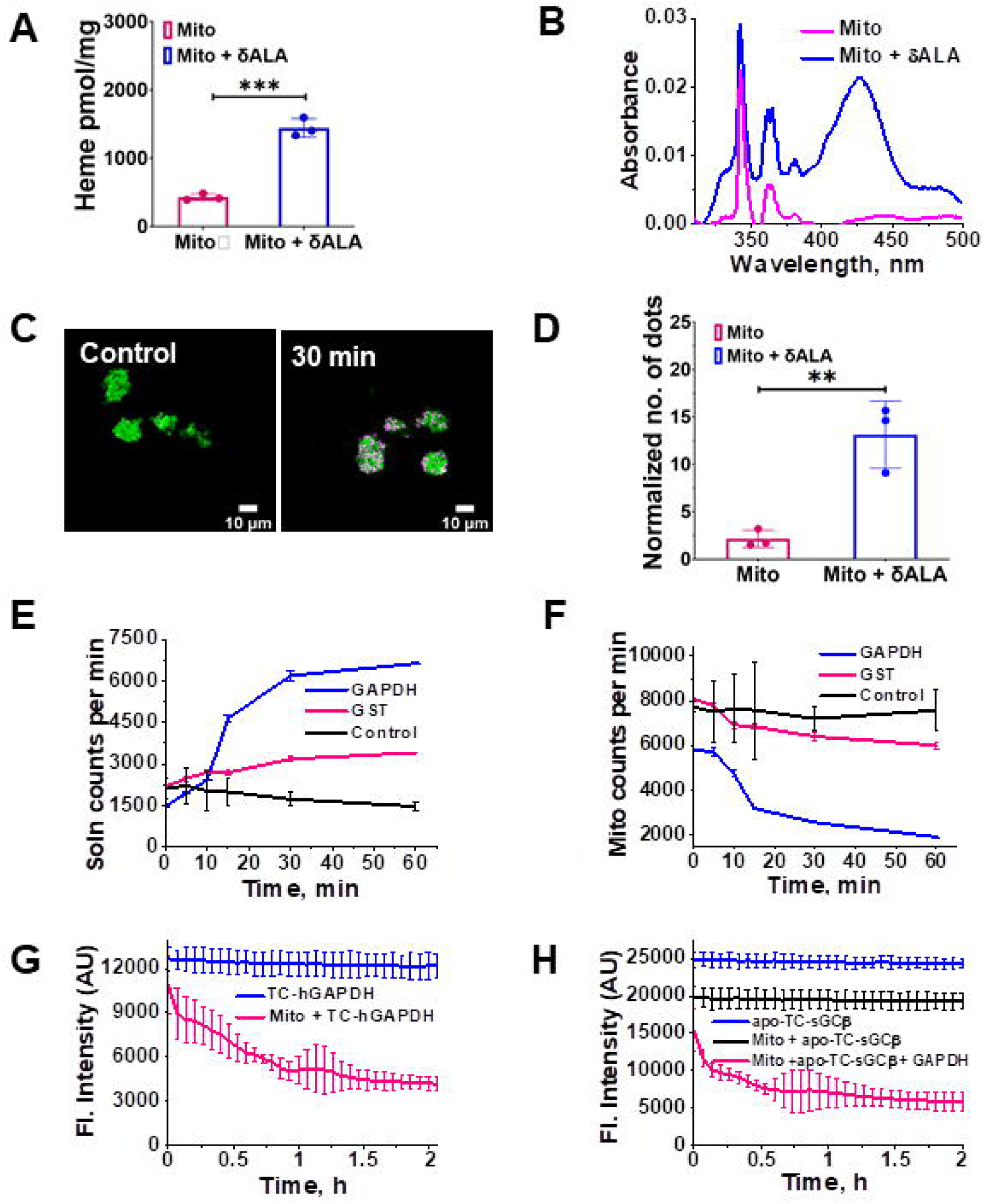
Heme export from isolated mitochondria is stimulated by GAPDH and involves direct FLVCR1b-GAPDH interaction and heme transfer. **A**, **B** Mitochondria that had been reisolated after being incubated *in vitro* for 30 min with δ-ALA and cell supernatant to trigger their heme biosynthesis contained an increased heme level upon their re-isolation as indicated by their total heme content and visible spectra. **C** Representative fluorescence microscope images of the mitochondria (HADHA, green) before or after the δ-ALA & cytosol incubation and re-isolation, indicating their relative levels of GAPDH-FLVCR1b interaction by PLA (red). **D** Quantification of the PLA results. **E**, **F** The impact of adding purified GAPDH or GST proteins to the isolated mitochondria on mitochondrial ^14^C-heme release into solution and loss of mitochondrial ^14^C-heme content, respectively. **G** Heme transfer to FlAsH-labeled TC-hGAPDH occurs upon addition of the isolated mitochondria at T = 0. **H** Adding purified GAPDH facilitates heme transfer from the isolated mitochondria to a target protein (FlAsH-apo-TC-sGCβ). **A, D, E, F**, Mean +/-SD of 3 independent trials. **G, H** Representative of three independent trials, mean +/- SD of triplicates. Significance: ** p < 0.01, *** p < 0.001 vs. the compared group based on a two-tailed t-test.

We then performed an experiment in which purified FlAsH-labeled TC-hGAPDH was added to the heme-loaded and reisolated mitochondria to determine if the observed release of heme caused by the GAPDH addition was tied to its binding of the released heme. Addition of TC-hGAPDH stimulated the release of mitochondrial heme as before, which bound to it as indicated by the time-dependent decrease in its fluorescence intensity (Fig. 3G). Finally, we examined if the GAPDH addition to the reisolated mitochondria would enable the released heme to be delivered to the GAPDH client protein TC-apo-sGCβ. As shown in Fig. 3H, the fluorescence of FlAsH-TC-apo-sGCβ when present alone in the reaction solution remained steady over a 1 h time course. Addition of the reisolated mitochondria to this solution caused an immediate partial decrease in the FlAsH-TC-apo-sGCβ fluorescence, which did not change with time and that we attribute to some build-up of heme that had become released from the mitochondrial preparation after re-isolation. Adding GAPDH to the reisolated mitochondria and FlAsH-TC-apo-sGCβ solution immediately stimulated additional heme to transfer into the FlAsH-TC-apo-sGCβ, as indicated by a time-dependent loss of its FlAsH fluorescence intensity (Fig. 3H). Thus, the main features of mitochondrial heme export to GAPDH and its subsequent delivery to a target protein that we had observed in living cells (*i.e.*, a time-dependent buildup of the FLVCR1b-GAPDH interaction coinciding with heme transfer to GAPDH and GAPDH heme transfer to the apo-sGCβ target protein) could be recapitulated in a simple reconstitution system consisting only of isolated mitochondria, purified GAPDH, and the purified apo-sGCβ target protein. In this system, GAPDH interacted with the mitochondrial FLVCR1b and had a strong stimulatory effect on the release of mitochondrial heme, which bound to the GAPDH and in turn allowed the heme to be delivered to the client protein.

### Assessing roles for the Progesterone Receptor Membrane Component 2 (PGRMC2) and the Transport and Golgi Organization 2 (TANGO2) proteins in FLVCR1b heme export to GAPDH and its heme allocation

Because two other proteins (PGRMC2 and TANGO2) have recently been implicated in intracellular heme trafficking ^28–31^ we tested for their involvement in our system. A siRNA knockdown of PGRMC2 expression in HEK293T cells reduced PGRMC2 expression by an average of 75% (Fig. S5A and B) but did not impact cell expression of other relevant proteins (Fig. S5C-E) nor did it inhibit mitochondrial heme transfer to TC-hGAPDH or inhibit the delivery of mitochondrial heme to downstream target proteins sGCβ or IDO1 in cells that were given δ-ALA/Fe (Fig. S6A-C). This indicated PGRMC2 was not involved and so it was not studied further. In contrast, a knockdown of TANGO2 expression in the HEK293T cells (Fig. S7A and B) strongly inhibited mitochondrial heme transfer to TC-hGAPDH following δ-ALA/Fe addition (Fig. 4A) and also blocked the downstream heme deliveries to sGCβ and to IDO1 (Fig. 4B and C), without impacting the expression levels of these proteins (Fig. S7C-E) or the cell heme level (Table S2). Thus, TANGO2 appeared to be involved in mitochondrial heme transfer to GAPDH, and we further investigated its role.

**Fig. 4:**
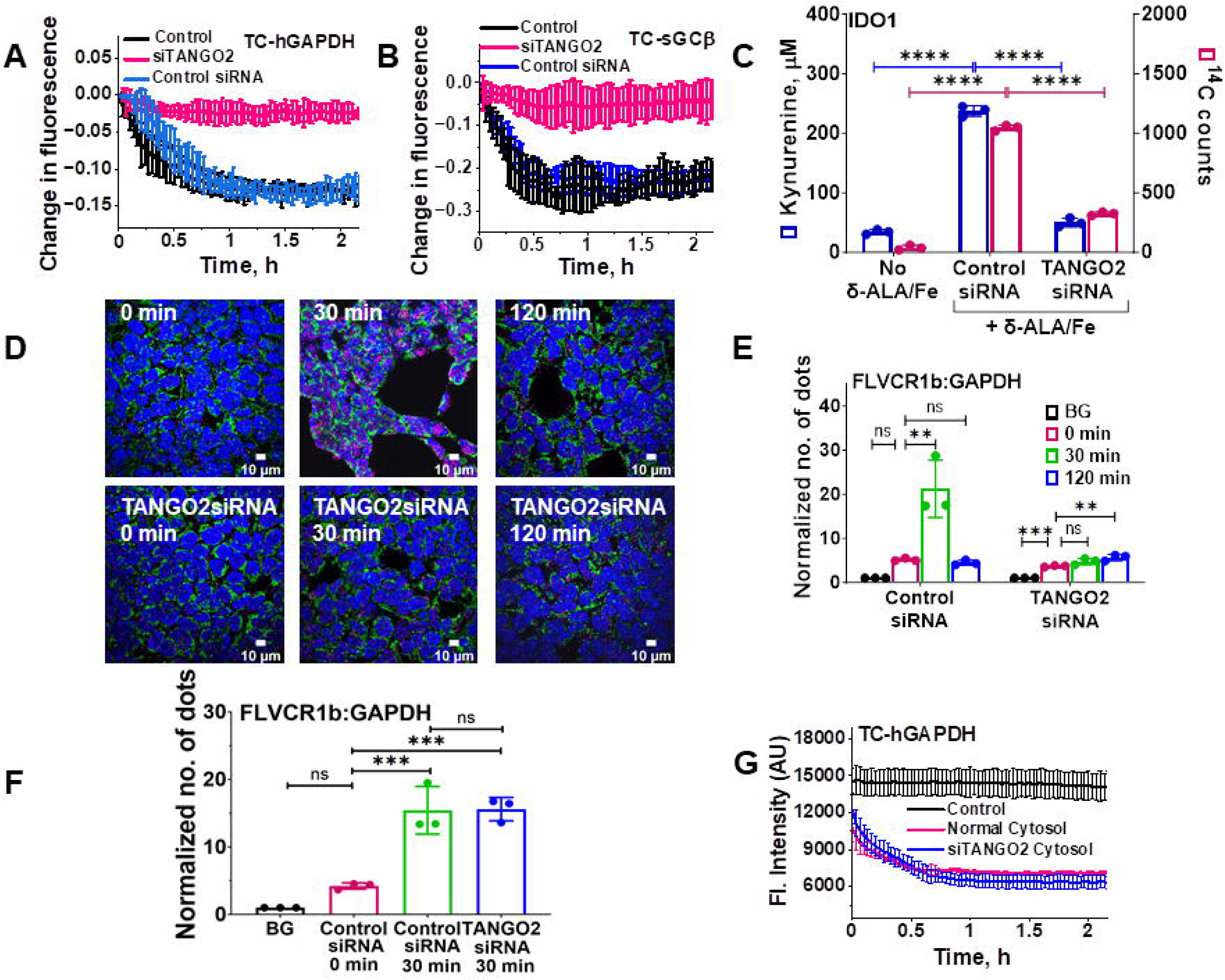
TANGO2 enables FLVCR1b heme export to GAPDH and consequent heme delivery to target proteins. **A** Targeted siRNA-based knockdown of cell TANGO2 expression inhibited mitochondrial heme transfer to FlAsH-labeled TC-hGAPDH in live HEK293T cells after their mitochondrial heme biosynthesis was stimulated by δ-ALA/Fe addition at time = 0. **B**, **C** TANGO2 knockdown inhibited the delivery of mitochondrial heme to two GAPDH client proteins in the living cells after the δ-ALA/Fe addition, as judged by **B** it blocking the decrease in fluorescence intensity for FlAsH-labeled TC-sGCβ and by **C** it blocking the gain in the enzymatic activity and ^14^C heme content of IDO1. **D** Representative fluorescence microscope images of cells stained with DAPI for nuclei (blue), using HADHA antibody for mitochondria (green), and the GAPDH-FLVCR1b interaction by PLA (red) in cells processed after the indicated times of δ-ALA/Fe exposure. BG, background. **E** Quantification of the PLA results to show the relative levels of GAPDH-FLVCR1b interaction. BG is the background PLA signal level. **F** PLA data comparing the relative interaction level of FLVCR1b and GAPDH on mitochondria that had been reisolated after having been incubated for the indicated times with δ-ALA plus supernatants prepared either from control or TANGO2 knockdown HEK293T cells. BG is the background PLA signal level. **G** Heme transfer from the reisolated mitochondria to FlAsH-labeled TC-hGAPDH after mitochondria were added at time = 0. **A, B, G** Representative of three independent trials, values are the mean +/- SD of triplicates. **C, E, F** Three independent trials, mean +/- SD. Significance: * p < 0.05, ** p < 0.01, *** p < 0.001, and **** p < 0.0001 vs. the compared group based on a one-way ANOVA test. ns, not significant.

Protein interaction studies using the PLA showed that knockdown of TANGO2 expression did not greatly alter the level of GAPDH interaction with FLVCR1b in resting cells, but it did prevent their interaction from increasing after δ-ALA/Fe was added (Fig. 4D & E). Further interrogation showed that TANGO2 interacted directly with FLVCR1b in the resting cells (5x above background) (Fig. S8A and B), and their level of interaction remained constant in cells that were given δ-ALA/Fe to increase mitochondrial heme production (Fig. S8A-C) and also did not change in cells that underwent a knockdown of GAPDH expression (Fig. S9A-C). In comparison, PLA results showed TANGO2 engaged in only a weak interaction with GAPDH (2.5x greater than background) both prior to or following δ-ALA/Fe addition to the cells (Fig. S10). Together, these results are consistent with TANGO2 primarily interacting with the FLVCR1b exporter to enable the mitochondrial heme transfer to GAPDH.

Because TANGO2 is thought to be localized to both cell cytosol and mitochondria ^32,33^, we further dissected its role by utilizing isolated mitochondria to examine the importance of cytosolic *versus* mitochondrial TANGO2 in enabling the mitochondrial heme transfer to TC-hGAPDH. We incubated isolated mitochondria with δ-ALA and with mitochondria-free cytosols that were prepared either from normal or from TANGO2 knockdown HEK293T cells, and then reisolated the mitochondria after the incubation and compared their extents of FLVCR1b-GAPDH interaction by PLA and their abilities to transfer heme to added TC-hGAPDH. The two sets of reisolated mitochondria behaved identically regarding their having an increased level of FLVCR1b-GAPDH interaction (Fig. 4F and S11) and in their ability to transfer heme to TC-hGAPDH (Fig. 4G). These results discount a role for cytosolic TANGO2 and instead implicate mitochondrial TANGO2 in enabling the FLVCR1b heme transfer to GAPDH.

## Discussion

How heme generated inside the mitochondria is released, distributed, and inserted into proteins that reside outside this organelle but require heme for function has been a long-standing unanswered question in cell biology ^1,4,5,34^. Our finding that the outer mitochondrial membrane transporter FLVCR1b exclusively provides heme to GAPDH finally completes an outline for the entire process. The unveiling of the key steps and players involved in eukaryotic heme allocation indicate the process is surprisingly simple and specific, and this information can now be leveraged to investigate the molecular details and regulation of this essential process. The results to date support the mechanism outlined in Fig. 5: Heme made in the mitochondria is provided to GAPDH in the cytosol solely through the FLVCR1b exporter. The heme transfer involves a direct protein-protein interaction between FLVCR1b and GAPDH, with their level of interaction being dynamic and changing according to mitochondrial heme availability and the extent of heme loading into GAPDH. After GAPDH obtains heme it dissociates from the mitochondria, which enables it to transport heme and make it available to numerous client proteins that are located in the cytosol or in other cell compartments. The target heme proteins are present in cells in their heme-free states and are typically are in complex with the cell chaperone Hsp90, which drives their heme insertions in an ATP-dependent manner, possibly with assistance from co-chaperone proteins ^13,35^. Hsp90 then dissociates from the heme-replete mature proteins, allowing their biological function ^18^. Regarding TANGO2, it had recently been proposed to act as a heme chaperone in cells or to aid in heme transfer from heme-enriched compartments, perhaps by acting independent of other heme-transporting proteins ^30,31^. Our current findings clarify its role by showing TANGO2 interacts with FLVCR1b and not with GAPDH and enables FLVCR1b export of mitochondrial heme to GAPDH. This role is consistent with knockout of TANGO2 causing an increase in the cell mitochondrial heme level ^31^, with TANGO2 displaying a poor affinity toward binding heme ^31^, and with a recent study that cast doubt on its functioning in intracellular heme transport ^36^.

**Fig. 5:**
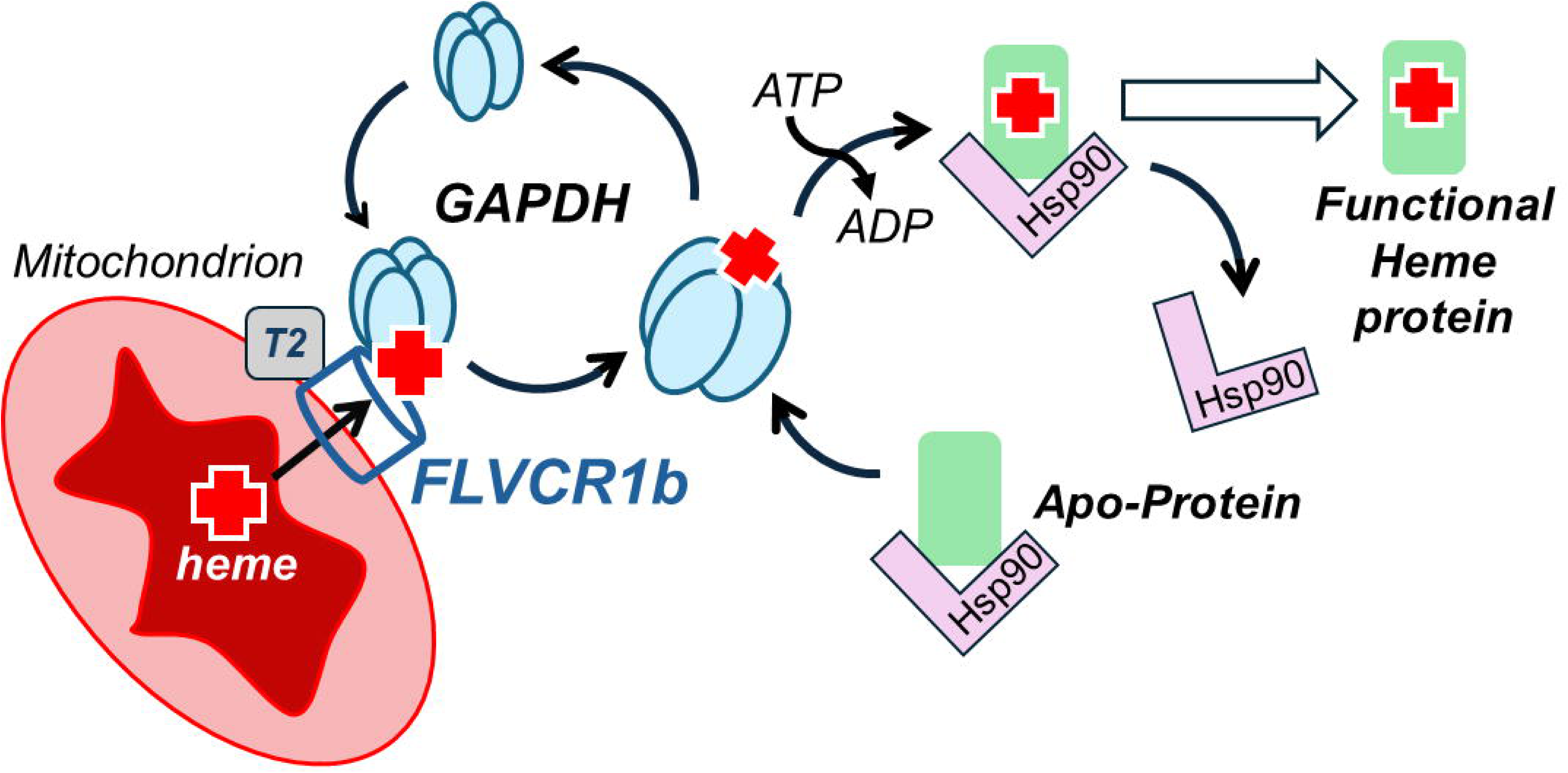
Pathway for mitochondrial heme provision in eukaryotic cells. Heme made in the mitochondria transfers to GAPDH solely through the mitochondrial FLVCR1b exporter. FLVCR1b functions in association with TANGO2 (T2), and its heme export involves a direct protein-protein interaction between FLVCR1b and GAPDH that is dynamic and influenced by the mitochondrial heme level and the extent of heme loaded onto the GAPDH. After GAPDH obtains heme, it dissociates from the mitochondria, and this makes heme broadly available to client proteins located in the cytosol or in other cell compartments. Client proteins are typically in complex with cell chaperone Hsp90, which is needed to drive their heme insertions in an ATP-dependent process. Hsp90 then dissociates, enabling heme protein biologic functions.

Given that there are other membrane transporters that may enable mitochondrial heme or porphyrin transport ^37,38^, and that membrane to membrane heme transfer has been invoked as a means for heme distribution in cells ^34,39^, it is surprising that mitochondrial heme provision to GAPDH and its downstream targets solely relies on the FLVCR1b transporter. This heralds a much broader role for FLVCR1b in enabling intracellular heme distribution, which prior to our study had only been shown to be required for the hemoglobinization that occurs during erythropoiesis ^20^. Indeed, the FLVCR1b-GAPDH heme transfer pathway that we describe can now explain why FLVCR1b is needed for hemoglobinization, because cellular heme delivery to apo-hemoglobin actually depends on GAPDH obtaining heme ^10^. In our current study, we also found that knockdown of FLVCR1b expression prevented delivery of mitochondrial heme to two other GAPDH-dependent client proteins (sGCβ and IDO1). This is consistent with FLVCR1b being the sole provider of mitochondrial heme to GAPDH, and with GAPDH being the sole heme source for these two downstream heme deliveries. Our findings can also explain why TANGO2 knockdown was found to prevent mitochondrial heme from reaching apo-myoglobin ^31^, because myoglobin acquisition of heme also depends on GAPDH obtaining heme in cells ^10^. Thus, without a functional FLVCR1b heme exporter to provide mitochondrial heme to GAPDH, it is unlikely that any GAPDH client protein will receive heme, thus severely compromising general heme allocation in cells. Indeed, one can envision GAPDH acting like a fountain that supplies mitochondrial heme to many proteins in the cell, and FLVCR1b acting as the only pipe that can supply the fountain with mitochondrial heme.

How FLVCR1b transfers mitochondrial heme to GAPDH should now be considered. FLVCR1b belongs to the major facilitator superfamily of transport proteins, which typically consist of 12 transmembrane-spanning helices connected by hydrophilic linkers that are exposed on either side of the membrane ^40,41^. These proteins fold into two domains consisting of transmembrane helices 1-6 and 7-12, forming an extensive interface between the domains that functions by a rocker mechanism to create alternating open channels on either side of the membrane that facilitate small ligand passage ^40^. FLVCR1b is the mitochondrially-targeted version of the larger cell membrane FLVCR1a transporter and is missing the first six transmembrane helices ^20^. Although the structures of FLVCR1a and b remain to be elucidated, FLVCR1b is expected to form a dimer in the membrane to enable the rocking mechanism for heme ligand passage ^42^. Within this context, it is fascinating that GAPDH stimulated heme export from the isolated mitochondria upon its addition. This behavior is reminiscent of how the addition of the heme-binding protein hemopexin increased heme flux out of cell membrane vesicles that contained the FLVCR1a exporter ^43^. Our findings indicate that GAPDH makes direct contact with FLVCR1b in cells and in isolated mitochondria, and imply their interaction mediates FLVCR1b heme export. Moreover, the GAPDH-FLVCR1b interaction increased as mitochondrial heme biosynthesis increased and then decreased in conjunction with heme loading into the GAPDH. This behavior may reflect structural changes in FLVCR1b and GAPDH. For example, an increase in mitochondrial heme availability might at first favor a conformational subpopulation of FLVCR1b that better interacts with GAPDH, followed by a subsequent change in GAPDH conformation upon its heme binding that diminishes their mutual affinity. Indeed, the affinity of GAPDH toward its client proteins is sensitive to their heme binding status ^18^. The basis for these changes in FLVCR1b-GAPDH interaction and how they may drive the heme transfer warrant further study.

## Conclusions

Our findings reveal that heme provision in eukaryotic cells relies on a surprisingly simple pathway that involves just four principal proteins: two of them (TANGO2 and FLVCR1b) enable mitochondrial heme export to one cytosolic protein (GAPDH) that then broadly provides it for distribution inside cells, and a fourth protein (Hsp90) that enables heme to become inserted into multiple protein clients. Together, this achieves a milestone in our understanding of heme biology and provides a foundation to investigate the cellular mechanisms that control heme allocation in eukaryotes and whether dysregulation occurs and underpins disease.

## Methods

### Reagents

All chemicals were purchased from Sigma unless otherwise noted. [^14^C]-δ-ALA (1 µCi/µl) was purchased from ChemDepo Inc. (Camarillo, CA). FlAsH-EDT_2_ was from Cayman Chemicals # 20704.

### Cell culture and expression of proteins by transfection

HEK293T cells (ATCC # CRL-3216) were cultured in Dulbecco’s Modified Eagle’s Medium (DMEM) (# 30-2002, ATCC) supplemented with 10% FBS (Gibco) at 37 °C and 5% carbon dioxide. At 60% confluence the cells were transfected using Lipofectamine 2000 (#11668019, Invitrogen) with various mammalian expression plasmids pRK5-TC-hGAPDH-HA or the H53A variant ^19^, pCMV5-TC-hsGCβ(1–619) ^44^, and pCMV3-hIDO1-FLAG (# HG11650-CF, Sino Biologicals). Protein expression was allowed for 48 h.

### Transfection of siRNA and gene silencing

Expression levels of select proteins in HEK293T cells were reduced using siRNA directed against human mRNAs of GAPDH (Dharmacon, #D-001830-02-20), FLVCR1b (Dharmacon, #CTM840033), PGRMC2 (Santa Cruz, #sc-88944), or TANGO2 (Sigma, #EHU064871). siRNA’s were used at a final concentration of 100 nM in the cell cultures along with Lipofectamine 2000. The siRNA-treated cells were cultured for 48 h before they received transfections with protein expression plasmids as described above.

### Cell supernatant preparation

HEK293T cells were harvested in using 50 mM Tris-HCl pH 7.4 buffer with 0.1% Triton X-100, 5 mM Na-molybdate and EDTA-free protease inhibitor cocktail (Roche). After three freeze-thaw cycles the lysate was centrifuged at 10,000 x g for 15 min at 4 °C and the supernatant was collected. Bradford (Bio-Rad # 500–0006) or DC method (Bio-Rad #5000111) were used to measure the protein concentration.

### Antibodies used for Western blot or immunoprecipitation

Human FLVCR (Novus Biologicals, #NB100-1481; dilution 1:1000), PGRMC2 (Santa Cruz, #sc-374624; dilution 1:1000), TANGO2 (Invitrogen, #PA5-51818, dilution 1:1000), GAPDH (Cell Signalling, #2118S; dilution 1:2000), HADHA (Santa Cruz, #sc-374497; dilution 1:1000), Guanylyl Cyclase β1 (ER-19) (Sigma #G4405; dilution 1:1000), and αβ-actin (Sigma #A1978; dilution 1:2500), and antibodies against HA (Cell Signaling, #3724S; dilution 1:2000) and FLAG (Sigma #F1804; dilution 1:1000).

### Western blot

Proteins were detected by chemiluminescence following Western blotting using HRP conjugated secondary antibodies of anti-mouse (Bio-Rad # 170–6516; dilution 1:10,000), anti-rabbit (GE Healthcare # NA9340; dilution 1:10,000) or anti-goat (Bio Rad #1721034; dilution 1:10,000) origin and ECL substrate (Thermo Scientific # 32106). Images were acquired and analyzed using a chemidoc system from Bio-Rad.

### Image analysis of Western blots

Image J quantification software (Image J; http://rsb.info.nih.gov/ij/) was used to quantify band intensities on western blots. Densitometric analysis (Image J; http://rsb.info.nih.gov/ij/) was used to measure relative protein amounts. The relative abundances of the protein of interest were determined by dividing its band intensity in each sample by the band intensity of a relevant protein control (i.e., FLVCR1b, actin) that was present and analyzed in the same sample.

### Measurement of total heme

Total heme in cell supernatants was measured by an established fluorometric method ^45^. Briefly, 20 µl of cell supernatant was mixed with 980 µl of 2 M oxalic acid and boiled for 1h. After cooling to room temperature and centrifugation, the total porphyrin was measured by its fluorescence emission at 662 nm (excitation 400 nm) relative to standard curves generated with freshly-prepared heme solutions.

### Isolation of cell cytoplasmic and mitochondrial fractions

To prepare mitochondria-free cytosol, HEK293T cells with and without TANGO2 silencing were lysed in cytosol extraction buffer (10 mM Tris-HCl, 0.34 mM Sucrose, 3 mM CaCl_2_, 2 mM MgCl_2_, 0.1 mM EDTA, 1 mM DTT, 5 mM Na-molybdate and EDTA-free protease inhibitor cocktail (Roche) on ice, then centrifuged at 720 x g for 5 min. The supernatant recovered was centrifuged at 10,000 x g for 5 min, and then was centrifuged 1 h in a Beckman Coulter ultracentrifuge (Optima L-100 XP) at 100,000 x g and then aliquoted and stored for use. Mitochondria were isolated from HEK293T cells at 4 °C following a previous procedure ^20^. Cells were lysed in mitochondrial extraction buffer (0.25 M sucrose, 10 mM HEPES pH 7.4, 5 mM Na-molybdate and EDTA-free protease inhibitor cocktail (Roche). The cell suspension was passed through a 1 mL syringe and 27 gauge needle 10 times. The homogenate was centrifuged twice at 600 x g for 5 min and then the supernatant was centrifuged at 10,000 x g for 10 min to pellet crude mitochondria. The crude mitochondrial pellet was diluted with mitochondrial extraction buffer to a 1 mg/ml protein concentration. For IP studies the mitochondrial suspension was then solubilized with 1% Triton X-100 and protease inhibitor on ice for 30 min, centrifuged at 12,000 x g for 10 min at 4 °C, and the supernatant used following protein quantification (Bio-Rad #5000111).

### Immunoprecipitation of mitochondrial samples

Solubilized mitochondrial supernatant samples (0.5 mg protein) were mixed with anti-FLVCR antibody (Novus Biologicals, #NB100-1481) and Protein G agarose beads (Cytiva # 17061801) to pull down the antibody-protein complex. The beads were washed three times with 1 mL lysis buffer (50 mM Tris-HCl pH 7.4 buffer with 0.1% Triton X-100, 5 mM Na-molybdate and EDTA-free protease inhibitor cocktail) by centrifugation (1000 x g for 5 min at 4°C). The beads were then boiled in Laemmli buffer, resolved onto 10% SDS-PAGE and Western transferred to PVDF membrane (Bio-Rad # 1620177) to probe for proteins of interest, using the antibodies listed above.

### Live cell heme binding kinetics using FIAsH-labeled TC-hGAPDH

HEK293T cells in black walled 96-well plates (Greiner Bio-One, # 655090) that had been grown in heme-deficient conditions, transfected to express TC-hGAPDH, and had or had not undergone FLVCR1b, PGRMC2, or TANGO2 silencing, were labelled with FlAsH using the method described previously ^19^. Briefly, the cell monolayers were washed once with phenol red-free Dulbecco’s modified Eagle’s medium (DMEM) containing 1 g/L glucose and then given FlAsH-EDT_2_ (5 μM) in Opti-MEM for 30 min at 37 °C. Afterward, the cell monolayers were washed twice with Phenol red free DMEM containing 10% heme depleted serum. Kinetic analyses were then performed in a FlexStation 3 plate reader (Molecular Devices) at 37 °C with excitation at 508 nm and emission at 528 nm. The fluorescence emission of FlAsH-labelled TC-hGAPDH in the cells was followed versus time with or without δ-ALA (1 mM) plus ferric citrate (100 µM) or hemin chloride alone (5 µM) being added to the cells at time = 0.

### Heme transfer kinetics into TC-sGCβ (1– 619) using FIAsH-EDT_2_

Heme transfer into FIAsH-labelled TC-sGCβ (1–619) expressed in HEK293T cells that had been grown in heme-deficient conditions and treated with corresponding siRNAs was monitored using methods reported previously ^44^ and as described above for TC-hGAPDH. Briefly, after FlAsH labelling and cell monolayer washing the cell fluorescence was monitored starting with or without addition of δ-ALA (1mM) and ferric citrate (100 µM) at time = 0. Kinetic analyses were then performed in a FlexStation 3 (Molecular Devices) plate reader at 37 °C with excitation at 508 nm and emission at 528 nm.

### Determination of ^14^C-labelled heme in IDO1

HEK293T cells were grown in DMEM and silenced for FLVCR1b, PGRMC2, or TANGO2 expression for 48 h before being transfected to express FLAG-IDO1. Transfected cells were then given ^14^C-δ-ALA (14 µM) and 10 µM ferric citrate and further cultured for 48 h. The cells were lysed using 50 mM Tris-HCl pH 7.4 buffer with 0.1% Triton X-100, 5 mM Na-molybdate, and EDTA-free protease inhibitor cocktail (Roche), and following centrifugation immunoprecipitations were done using supernatant (I mg protein) and anti-FLAG antibody (Sigma #F1804; dilution 1:1000) and Protein G agarose beads (60 µL, Cytiva # 17061801) were used to pull down the antibody–protein complex. The beads were washed and isolated as described above and tubes containing the beads were inserted into scintillation vials which received 4 ml of scintillation fluid (Liquiscint, National Diagnostics # LS-121), and ^14^C counts were recorded with a scintillation counter as described ^12^.

### IDO1 activity assay

IDO1 activity was determined by colorimetric measure of the accumulated L-Kynurenine product in the cell culture medium. Cells were given L-Trp (2 mM) in phenol red free DMEM containing 10% FBS with or without δ-ALA (1 mM) and ferric citrate (100 µM). After 5 h of incubation the culture medium was de-proteinized by adding an equal volume of 3% trichloroacetic acid (Sigma # T6399) and incubation at 50 °C for 30 min. After centrifugation at 9000 x g for 10 min the supernatants were mixed with an equal volume of p-dimethyl-aminobenzaldehyde (Sigma # 109762; 20 mg/ml) in glacial acetic acid at room temperature and incubated 3 min to allow for chromophore formation. Absorbance at 492 nm was measured in a plate reader (Molecular Devices) and similarly-processed standards containing L-Kyn (Sigma #K8625) were used to calculate the sample L-Kyn concentrations.

### Mitochondrial preparation from mouse brain and liver

Mitochondria were isolated from brain or liver of 3-4 weeks old mice (18-22 g). The procedures were approved by Institutional Animal Care and Use Committee (IACUC) of the Cleveland Clinic. Mice were sacrificed by CO_2_ asphyxiation and cervical dislocation. The brain and liver were removed immediately and immersed in ice cold mitochondrial isolation buffer (MIB, 0.25 M sucrose, 0.5mM EDTA, 10 mM Tris-HCl, pH 7.4), and mitochondria were isolated by a discontinuous Percoll gradient method at 4°C as described previously ^46^. Briefly, Percoll (Sigma Aldrich #P1644) was diluted with MIB to obtain final concentrations of 12%, 26%, and 40% (v/v). Dounce homogenizers with glass pestles were used to homogenize each brain or liver sample in 12% Percoll. A gradient was prepared by pouring 26% Percoll on 40% Percoll. The homogenate in 12% Percoll was layered onto the gradient and was centrifuged at 30,000 x g in a Beckman Coulter ultracentrifuge (Optima L-100 XP) for 5 min. The second fraction was collected after removing the top layer and diluted 1:4 in MIB followed by centrifugation at 15,000 x g for 10 min. The pellet was resuspended in 1 ml of MIB and centrifuged at 15,000 x g for 5 min. The final mitochondrial pellet was resuspended in 100 µl of MIB, its protein concentration measured, and it was then used immediately for studies.

### Mitochondrial heme biosynthesis and heme labelling with ^14^C

Mitochondrial heme production was stimulated by resuspending mitochondria (5 mg protein) and mitochondria-free HEK293T cell cytosol (5 mg protein) to a final volume of 1 mL with incubation buffer (80 mM KCl, 50 mM MOPS, 5mM KH_2_PO_4_, 1mM EGTA, pH 7.4). This had δ-ALA (1mM) and 4 µM ADP (Sigma #A2754) added and was then incubated for 30 min at 37° C. In some cases, mitochondria-free cytosol from si-TANGO2 treated HEK293T cells was used, and in other cases 10 µM of ^14^C-δ-ALA (0.5 μCi/mg) was used. At the end of the incubations, mitochondria were pelleted by centrifugation (10,000 x g), washed twice, and then resuspended in incubation buffer (80 mM KC1, 50 mM MOPS, 5 mM K-phosphate, 1 mM EGTA, pH 7.4). Their heme content was quantified using the heme chromogen assay ^45^. Briefly, 90 µl of mitochondrial suspension was mixed with 160 µl of heme chromogen reagent (40:60 pyridine:H_2_O, 200 mM NaOH), the heme iron was reduced by adding a few grains of sodium dithionite, the absorbance was measured in a 96 well plate at 556 nm, and was quantified by standard curve based on freshly made heme samples.

### 14C-heme efflux from purified mitochondria

Heme efflux from the mitochondria isolated from mouse liver and brain was studied at room temperature according to a previously reported procedure ^25^. The ^14^C -heme-containing mitochondria (5 mg protein) prepared as above were incubated with or without 5 µM ADP and 20 µM of GAPDH or GST proteins for different times (0, 5, 10, 15, 30, 60 min). Reactions were stopped at each time point by transfer to ice and the samples were then centrifuged at 4 °C at 10,000 x g for 5 min. The amount of ^14^C-heme present in each sample supernatant and pellet was determined by scintillation counting as described previously ^12^.

### Visualizing heme efflux from mitochondria to TC-hGAPDH

Wells containing FIAsH-labelled TC-hGAPDH (2 µM) and ADP (0.5 µM) in a black-walled 96-well plate at room temperature had added to them either buffer alone or containing mitochondria (0.5 mg protein) that had been reisolated after being pre-incubated with δ-ALA and cytosols generated from either normal or TANGO2 knockdown HEK293T cells as described above. Fluorescence monitoring commenced immediately in a SYNERGY H1 microplate reader (BioTek) at room temperature with excitation at 508 nm and emission collection at 528 nm.

### Visualizing heme efflux from mitochondria to apo-TC-sGCβ

Wells in a black-walled 96-well plate containing FIAsH-labelled apo-TC-sGCβ (2 µM) and ADP (0.5 µM) at room temperature had added to them either buffer alone, buffer plus heme-loaded mitochondria (0.5 mg protein) prepared as above, or buffer containing the mitochondria and rabbit GAPDH (5 µM). Fluorescence monitoring commenced immediately in a SYNERGY H1 microplate reader (BioTek) at room temperature with excitation at 508 nm and emission collection at 528 nm.

### Proximity ligation assay (PLA) of protein-protein interactions

PLA was carried out according to the manufacturer’s protocol using Duolink In Situ Detection Reagents (Sigma Aldrich, #DUO92013) to visualize the intracellular protein-protein interactions. HEK293T cells were seeded onto glass coverslips (EMS # 7220010) immersed in 6 well plates (Genesee Scientific, # 25105) and cultured with SA (400 μM) and heme-depleted serum for 2 days to deplete cell heme stores. The coverslips were then washed and incubated with cell culture media plus δ-ALA (1mM) and Ferric Citrate (100 µM). Coverslips were removed at different times (0, 15, 30, 45, 60 and 120 min) and were immediately fixed with 4% formaldehyde (Sigma, #F-1268), washed 3x with PBS for 5min, and then the cells were permeabilized with 0.3% Triton X-100. Afterward, blocking was done (Sigma Aldrich, #DUO82007) followed by overnight incubations with various pairs of primary antibodies directed either against FLVCR1 (Novus Biologicals, #NB100-1481, dilution 1:100), TANGO2 (Invitrogen, #PA5-51818, dilution 1:100), GAPDH (Cell Signalling, #2118S, dilution 1:400), HADHA (Santa Cruz, #sc-374497, dilution 1:100), or HA (Cell Signalling, #3724S; dilution 1:400). After overnight incubation the primary antibodies were washed away and PLA secondary probes were added: Anti-Rabbit MINUS (Sigma Aldrich, #DUO92005, dilution 1:5), Anti-Goat PLUS (Sigma Aldrich, #DUO92003, dilution 1:5) and AlexaFluor 488 anti-Mouse secondary antibody (ThermoFischer, #A-21202, dilution 1:1000), followed by 1 h incubation in a humidified chamber at 37 °C. The coverslips were twice washed in 1x Duolink In Situ wash buffer A (Sigma Aldrich, #DUO82046) for 5 min under gentle agitation. Ligation was then done by adding Ligation buffer (Sigma Aldrich, #DUO82009, dilution 1:5) and Ligase (Sigma Aldrich, #DUO82027, dilution 1:40) solutions to each sample and a further 30 min incubation in a humidified chamber at 37 °C. The coverslips were then washed twice in 1x Duolink In Situ wash buffer A for 2 min under gentle agitation, and then were placed in amplification buffer (Sigma Aldrich, #DUO82012, dilution 1:5) and polymerase (Sigma Aldrich, #DUO82028, dilution 1:80) solution and incubated 100 min at 37 °C, followed by washing twice for 10 min in 1x wash buffer B (Sigma Aldrich, #DUO82048) and then once for 1 min in 100 x diluted wash buffer B, and drying at room temperature in the dark. Coverslips were individually mounted with Duolink In Situ Mounting Medium with DAPI (Sigma Aldrich, #DUO82040) onto slides, were sealed with nail polish, and were imaged under a confocal microscope (Inverted Leica SP8 confocal microscope) with an objective 63x. To determine the background PLA signal, the same procedures were followed as described above except that the cell-containing coverslips were incubated with only one of the individual primary antibodies instead of with a pair.

In some cases, the same PLA procedures were done for samples of mitochondria that had undergone incubation for different times with δ-ALA and cytosol, reisolation by centrifugation twice at 10,000 x g, and placement onto poly-D-lysine coated coverslips. Microscope images for mitochondrial samples utilized a 100x objective.

### Image analysis of PLA data

In the Duolink PLA, protein-protein interactions and are visualized as red dots ^24^. The numbers of dots per nuclei in each image were quantified by Volocity 6.5.1 (Quorum Technologies) image analysis software. Using the compartmentalization method, two populations (Blue DAPI for nuclei and magenta dots for protein-protein interaction) were selected and the populations in each compartment were calculated in the software. The number of dots per nuclei were averaged from each image. For each condition, experiments were done in three independent trials. Images used to determine the background PLA signal for each experimental circumstance were obtained from experiments as described above and were similarly analyzed to determine the average number of background dots per nuclei, which was reported as a relative value of 1.0 in the data presentations. The imaging results are presented as the mean of the three trial values ± standard deviation. Statistical significance (p-values) were determined using a one-way ANOVA test in the software GraphPad Prism (v9).

### Statistical analyses

The results are presented as the mean of the three trial values ± standard deviation. GraphPad Prism (v9) was used for the statistical analysis to measure significance (p-values) by one-way ANOVA or a two-tailed Student’s t-test.

## Funding and additional information

This work was supported by National Institutes of Health grants R01 GM130624 and R01 GM148664 (to D.J. Stuehr). The content is solely the responsibility of the authors and does not necessarily represent the official views of the National Institutes of Health.

## Acknowledgments

We thank all members of the Stuehr laboratory for helpful discussion and Dr. Nayden Naydenov (Cleveland Clinic) for providing us with mouse tissues for this study.

## Abbreviations

ATP: Adenosine triphosphate
FlAsH-EDT2: Fluorescein arsenical hairpin binder-ethane dithiol
GAPDH: Glyceraldehyde 3-phosphate dehydrogenase
HA: Hemagglutinin
Hsp90: Heat shock protein 90
IDO1: Indoleamine 2,3-dioxygenase 1
SA: Succinyl acetone
sGC: Soluble guanylyl cyclase
TC-hGAPDH: Tetra cysteine-human glyceraldehyde 3-phosphate dehydrogenase
δ-ALA: delta-aminolevulinic acid
FLVCR1b: Feline leukemia virus subgroup C receptor 1 isoform b
TANGO2: Transport and Golgi organization 2 homolog
PGRMC2: Progesterone receptor membrane component 2.

## References

1 Swenson, S. A. et al. From Synthesis to Utilization: The Ins and Outs of Mitochondrial Heme. Cells 9, doi:10.3390/cells9030579 (2020).

2 Reddi, A. R. & Hamza, I. Heme Mobilization in Animals: A Metallolipid’s Journey. Acc Chem Res 49, 1104–1110, doi:10.1021/acs.accounts.5b00553 (2016).

3 Dailey, H. A. & Medlock, A. E. A primer on heme biosynthesis. Biol Chem 403, 985–1003, doi:10.1515/hsz-2022-0205 (2022).

4 Gallio, A. E., Fung, S. S., Cammack-Najera, A., Hudson, A. J. & Raven, E. L. Understanding the Logistics for the Distribution of Heme in Cells. JACS Au 1, 1541–1555, doi:10.1021/jacsau.1c00288 (2021).

5 Ponka, P. Cell biology of heme. Am J Med Sci 318, 241–256, doi:10.1097/00000441-199910000-00004 (1999).

6 Hannibal, L. et al. Heme binding properties of glyceraldehyde-3-phosphate dehydrogenase. Biochemistry 51, 8514–8529, doi:10.1021/bi300863a (2012).

7 Sirover, M. A. The role of posttranslational modification in moonlighting glyceraldehyde-3-phosphate dehydrogenase structure and function. Amino Acids 53, 507–515, doi:10.1007/s00726-021-02959-z (2021).

8 White, M. R. & Garcin, E. D. D-Glyceraldehyde-3-Phosphate Dehydrogenase Structure and Function. Subcell Biochem 83, 413–453, doi:10.1007/978-3-319-46503-6_15 (2017).

9 Fleischhacker, A. S. & Ragsdale, S. W. An unlikely heme chaperone confirmed at last. J Biol Chem 293, 14569–14570, doi:10.1074/jbc.H118.005247 (2018).

10 Tupta, B. et al. GAPDH is involved in the heme-maturation of myoglobin and hemoglobin. FASEB J 36, e22099, doi:10.1096/fj.202101237RR (2022).

11 Chakravarti, R., Aulak, K. S., Fox, P. L. & Stuehr, D. J. GAPDH regulates cellular heme insertion into inducible nitric oxide synthase. Proc Natl Acad Sci U S A 107, 18004–18009, doi:10.1073/pnas.1008133107 (2010).

12 Sweeny, E. A. et al. Glyceraldehyde-3-phosphate dehydrogenase is a chaperone that allocates labile heme in cells. J Biol Chem 293, 14557–14568, doi:10.1074/jbc.RA118.004169 (2018).

13 Morishima, Y., Lau, M., Pratt, W. B. & Osawa, Y. Dynamic cycling with a unique Hsp90/Hsp70-dependent chaperone machinery and GAPDH is needed for heme insertion and activation of neuronal NO synthase. J Biol Chem 299, 102856, doi:10.1016/j.jbc.2022.102856 (2023).

14 Dai, Y., Sweeny, E. A., Schlanger, S., Ghosh, A. & Stuehr, D. J. GAPDH delivers heme to soluble guanylyl cyclase. J Biol Chem 295, 8145–8154, doi:10.1074/jbc.RA120.013802 (2020).

15 Islam, S., Jayaram, D. T., Biswas, P. & Stuehr, D. J. Functional maturation of cytochromes P450 3A4 and 2D6 relies on GAPDH- and Hsp90-Dependent heme allocation. J Biol Chem 300, 105633, doi:10.1016/j.jbc.2024.105633 (2024).

16 Dai, Y. et al. Heme delivery to heme oxygenase-2 involves glyceraldehyde-3-phosphate dehydrogenase. Biol Chem 403, 1043–1053, doi:10.1515/hsz-2022-0230 (2022).

17 Biswas, P., Dai, Y. & Stuehr, D. J. Indoleamine dioxygenase and tryptophan dioxygenase activities are regulated through GAPDH- and Hsp90-dependent control of their heme levels. Free Radic Biol Med 180, 179–190, doi:10.1016/j.freeradbiomed.2022.01.008 (2022).

18 Stuehr, D. J., Dai, Y., Biswas, P., Sweeny, E. A. & Ghosh, A. New roles for GAPDH, Hsp90, and NO in regulating heme allocation and hemeprotein function in mammals. Biol Chem 403, 1005–1015, doi:10.1515/hsz-2022-0197 (2022).

19 Biswas, P. et al. Visualizing mitochondrial heme flow through GAPDH in living cells and its regulation by NO. Redox Biol 71, 103120, doi:10.1016/j.redox.2024.103120 (2024).

20 Chiabrando, D. et al. The mitochondrial heme exporter FLVCR1b mediates erythroid differentiation. J Clin Invest 122, 4569–4579, doi:10.1172/JCI62422 (2012).

21 Adams, S. R. et al. New biarsenical ligands and tetracysteine motifs for protein labeling in vitro and in vivo: synthesis and biological applications. J Am Chem Soc 124, 6063–6076, doi:10.1021/ja017687n (2002).

22 Ito, H. et al. Oral administration of 5-aminolevulinic acid induces heme oxygenase-1 expression in peripheral blood mononuclear cells of healthy human subjects in combination with ferrous iron. Eur J Pharmacol 833, 25–33, doi:10.1016/j.ejphar.2018.05.009 (2018).

23 Doty, R. T., Sanchez-Bonilla, M., Keel, S. B. & Abkowitz, J. L. FLVCR1a But Not FLVCR1b Is Required For Effective Erythropoiesis In Adult Mice. Blood 122, 308–308, doi:10.1182/blood.V122.21.308.308 (2013).

24 Alam, M. S. Proximity Ligation Assay (PLA). Curr Protoc Immunol 123, e58, doi:10.1002/cpim.58 (2018).

25 Liem, H. H., Grasso, J. A., Vincent, S. H. & Muller-Eberhard, U. Protein-mediated efflux of heme from isolated rat liver mitochondria. Biochem Biophys Res Commun 167, 528–534, doi:10.1016/0006-291x(90)92056-6 (1990).

26 Harvey, J. W. & Beutler, E. Binding of heme by glutathione S-transferase: a possible role of the erythrocyte enzyme. Blood 60, 1227–1230 (1982).

27 Senjo, M., Ishibashi, T. & Imai, Y. Purification and characterization of cytosolic liver protein facilitating heme transport into apocytochrome b5 from mitochondria. Evidence for identifying the heme transfer protein as belonging to a group of glutathione S-transferases. J Biol Chem 260, 9191–9196 (1985).

28 Chen, C. & Hamza, I. Notes from the Underground: Heme Homeostasis in C. elegans. Biomolecules 13, doi:10.3390/biom13071149 (2023).

29 Galmozzi, A. et al. PGRMC2 is an intracellular haem chaperone critical for adipocyte function. Nature 576, 138–142, doi:10.1038/s41586-019-1774-2 (2019).

30 Han, S., Guo, K., Wang, W., Tao, Y. J. & Gao, H. Bacterial TANGO2 homologs are heme-trafficking proteins that facilitate biosynthesis of cytochromes c. mBio 14, e0132023, doi:10.1128/mbio.01320-23 (2023).

31 Sun, F. et al. HRG-9 homologues regulate haem trafficking from haem-enriched compartments. Nature 610, 768–774, doi:10.1038/s41586-022-05347-z (2022).

32 Jennions, E. et al. TANGO2 deficiency as a cause of neurodevelopmental delay with indirect effects on mitochondrial energy metabolism. J Inherit Metab Dis 42, 898–908, doi:10.1002/jimd.12149 (2019).

33 Milev, M. P. et al. The phenotype associated with variants in TANGO2 may be explained by a dual role of the protein in ER-to-Golgi transport and at the mitochondria. J Inherit Metab Dis 44, 426–437, doi:10.1002/jimd.12312 (2021).

34 Chambers, I. G., Willoughby, M. M., Hamza, I. & Reddi, A. R. One ring to bring them all and in the darkness bind them: The trafficking of heme without deliverers. Biochim Biophys Acta Mol Cell Res 1868, 118881, doi:10.1016/j.bbamcr.2020.118881 (2021).

35 Ghosh, A., Dai, Y., Biswas, P. & Stuehr, D. J. Myoglobin maturation is driven by the hsp90 chaperone machinery and by soluble guanylyl cyclase. FASEB J 33, 9885–9896, doi:10.1096/fj.201802793RR (2019).

36 Sandkuhler, S. E. et al. Haem’s relevance genuine? Re-visiting the roles of TANGO2 homologues including HRG-9 and HRG-10 in C. elegans. bioRxiv, doi:10.1101/2023.11.29.569072 (2023).

37 Donegan, R. K., Moore, C. M., Hanna, D. A. & Reddi, A. R. Handling heme: The mechanisms underlying the movement of heme within and between cells. Free Radic Biol Med 133, 88–100, doi:10.1016/j.freeradbiomed.2018.08.005 (2019).

38 Schaedler, T. A. et al. Structures and functions of mitochondrial ABC transporters. Biochem Soc Trans 43, 943–951, doi:10.1042/BST20150118 (2015).

39 Rose, M. Y., Thompson, R. A., Light, W. R. & Olson, J. S. Heme transfer between phospholipid membranes and uptake by apohemoglobin. J Biol Chem 260, 6632–6640 (1985).

40 Law, C. J., Maloney, P. C. & Wang, D. N. Ins and outs of major facilitator superfamily antiporters. Annu Rev Microbiol 62, 289–305, doi:10.1146/annurev.micro.61.080706.093329 (2008).

41 Tailor, C. S., Willett, B. J. & Kabat, D. A putative cell surface receptor for anemia-inducing feline leukemia virus subgroup C is a member of a transporter superfamily. J Virol 73, 6500–6505, doi:10.1128/JVI.73.8.6500-6505.1999 (1999).

42 Chiabrando, D. et al. Expression and purification of the heme exporter FLVCR1a. Protein Expr Purif 172, 105637, doi:10.1016/j.pep.2020.105637 (2020).

43 Yang, Z. et al. Kinetics and specificity of feline leukemia virus subgroup C receptor (FLVCR) export function and its dependence on hemopexin. J Biol Chem 285, 28874–28882, doi:10.1074/jbc.M110.119131 (2010).

44 Dai, Y., Faul, E. M., Ghosh, A. & Stuehr, D. J. NO rapidly mobilizes cellular heme to trigger assembly of its own receptor. Proc Natl Acad Sci U S A 119, doi:10.1073/pnas.2115774119 (2022).

45 Albakri, Q. A. & Stuehr, D. J. Intracellular assembly of inducible NO synthase is limited by nitric oxide-mediated changes in heme insertion and availability. J Biol Chem 271, 5414–5421, doi:10.1074/jbc.271.10.5414 (1996).

46 Rajapakse, N., Shimizu, K., Payne, M. & Busija, D. Isolation and characterization of intact mitochondria from neonatal rat brain. Brain Res Brain Res Protoc 8, 176–183, doi:10.1016/s1385-299x(01)00108-8 (2001).

