## Supplemental Data for "Heme allocation in eukaryotic cells relies on mitochondrial heme export through FLVCR1b to cytosolic GAPDH"

Table S1. Total soluble heme content in control and si-RNA treated HEK293T cell supernatants

Values are the mean +/- SD from 3 independent experiments

| Control (pmol/mg protein) | siFLVCR1b (pmol/mg protein) |
|---|---|
| 65 ± 5 | 60 ± 4 |

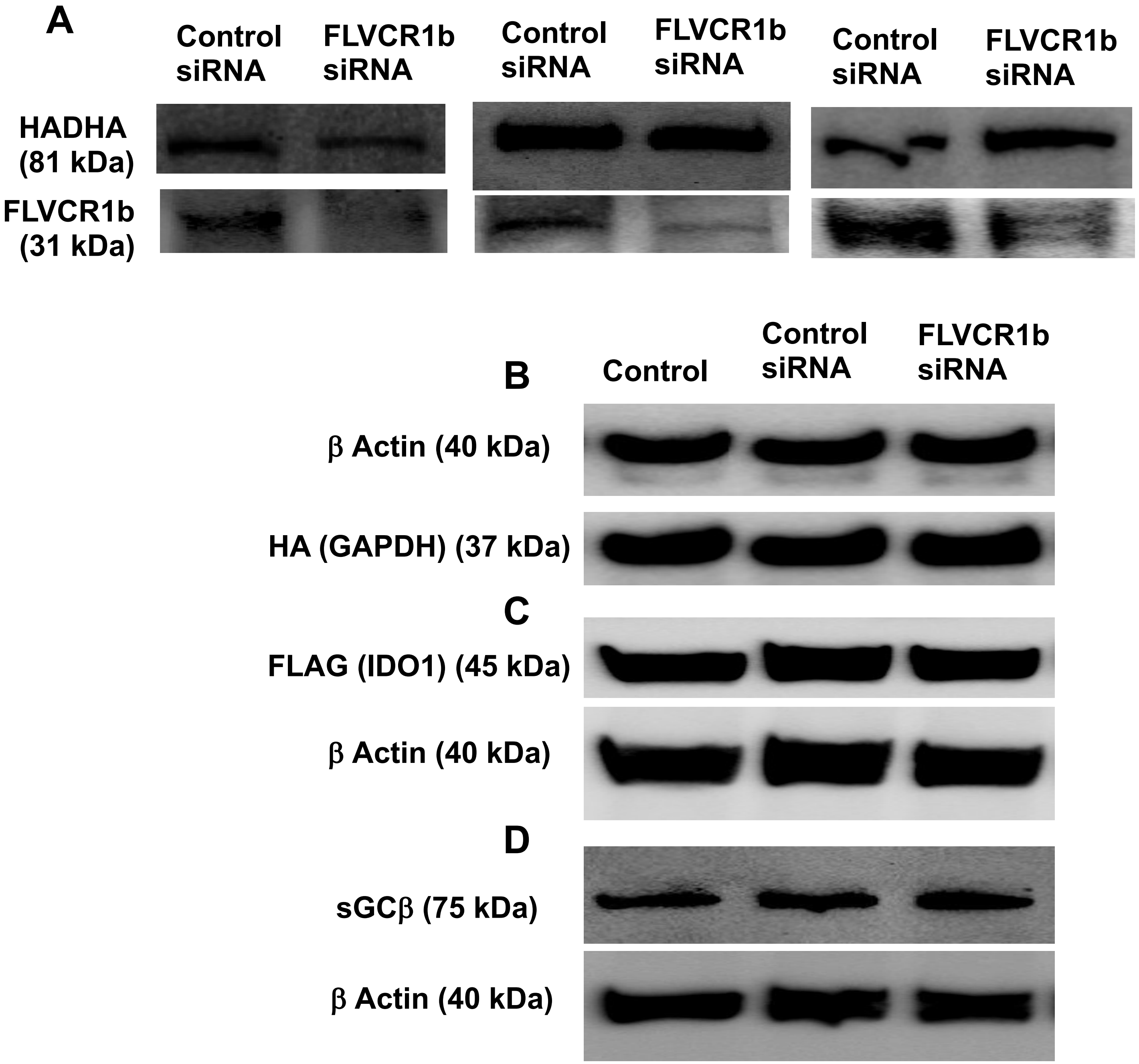


**Supplementary Figure 1. Impact of scrambled (control) or FLVCR1b-targeted siRNA treatment on the expression levels of the indicated proteins in transfected HEK293T cells.** The samples used here were the same as those reported on in main Figure 1, and the images are from representative Western blots. Equal total protein amounts of each cell supernatant sample were run on SDS-PAGE and Western blotted and developed using antibodies against the indicated proteins or tags. **A** HADHA and FLVCR1b expression levels in response to the scrambled or targeted siRNA treatment. **B, C, D** Expression levels of HA-GAPDH, FLAG-IDO1, and sGCβ relative to β-actin expression and in response to the scrambled or targeted siRNA treatment.

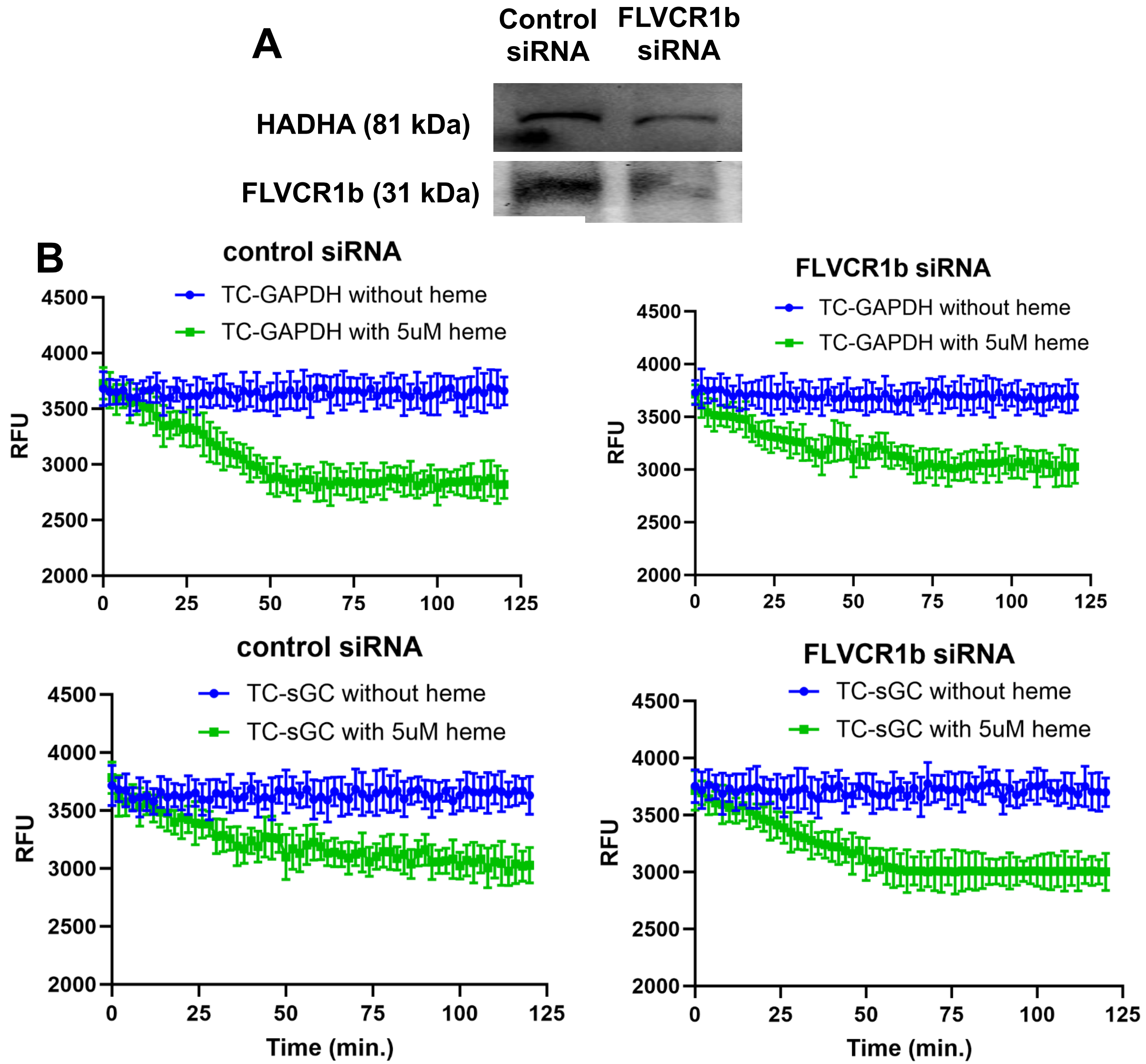


**Supplementary Figure 2. Effect of FLVCR1b knockdown on the ability of TC-GAPDH to bind exogenous heme provided to the cell cultures.** HEK293T cells expressing TC-hGAPDH underwent scrambled (control) or targeted siRNA treatment and then had vehicle or vehicle + heme added into the culture media. Heme binding by FlAsH-labeled TC-hGAPDH expressed in the cells was followed by monitoring their fluorescence versus time. **A** Representative Western blot comparing levels of HADHA or FLVCR1b expression in cells that underwent scrambled or targeted siRNA treatments. **B** Fluorescence intensity versus time traces (data points are mean +/- SD, n = 3 wells) recorded after the cell cultures received vehicle or vehicle plus heme. Panels show results from two independent trials.

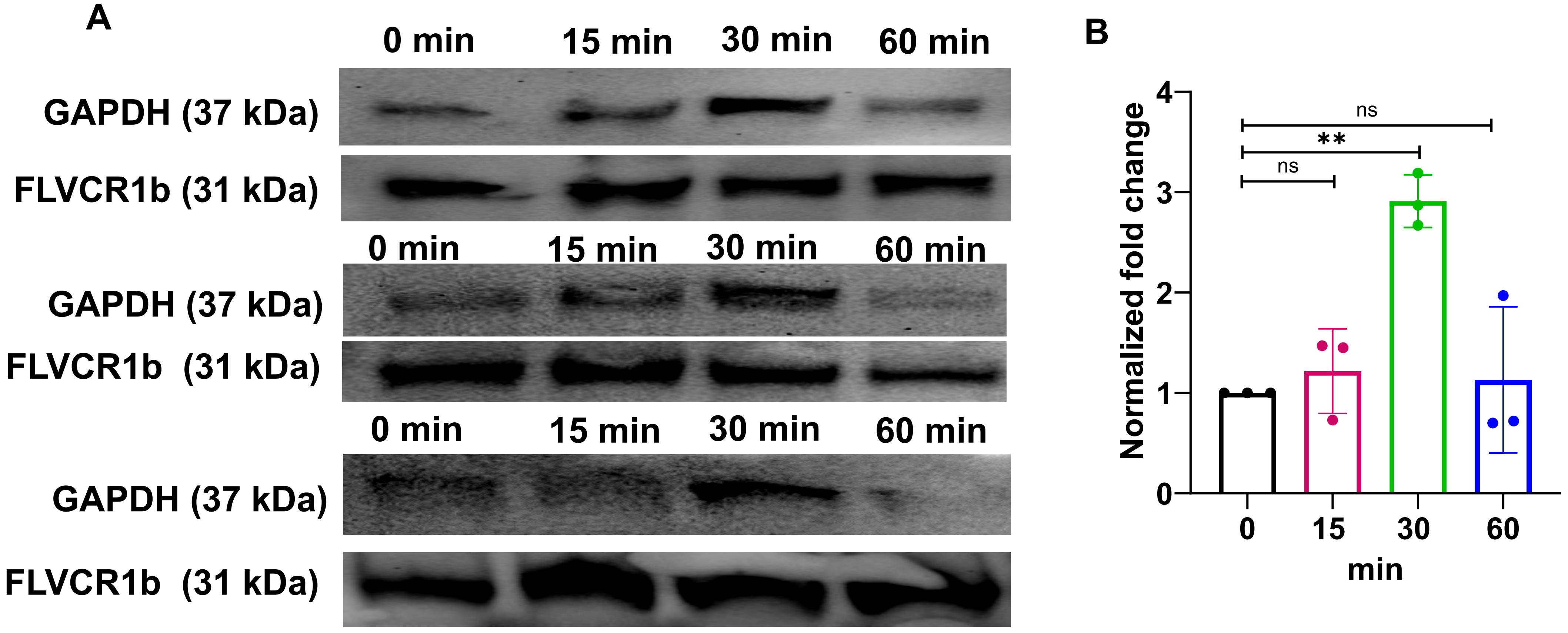


**Supplementary Figure 3. Time course of FLVCR1b interaction with GAPDH in cells after stimulation of their mitochondrial heme biosynthesis.** Supernatant samples (equal total protein) from HEK293T cells that were harvested at the indicated time points after δ-ALA/Fe addition were subject to pulldown using anti-FLVCR1b antibody. **A** Representative Western blots showing relative levels of FLVCR1b and GAPDH proteins in the pulldown samples. **B** Fold change in the GAPDH interaction with FLVCR1b versus time after the δ-ALA/Fe addition as determined from quantitation of the GAPDH Western band intensities. Three independent trials, mean +/- SD. Significance: ** $p < 0.01$ vs. the compared group based on a one-way ANOVA. ns, not significant.

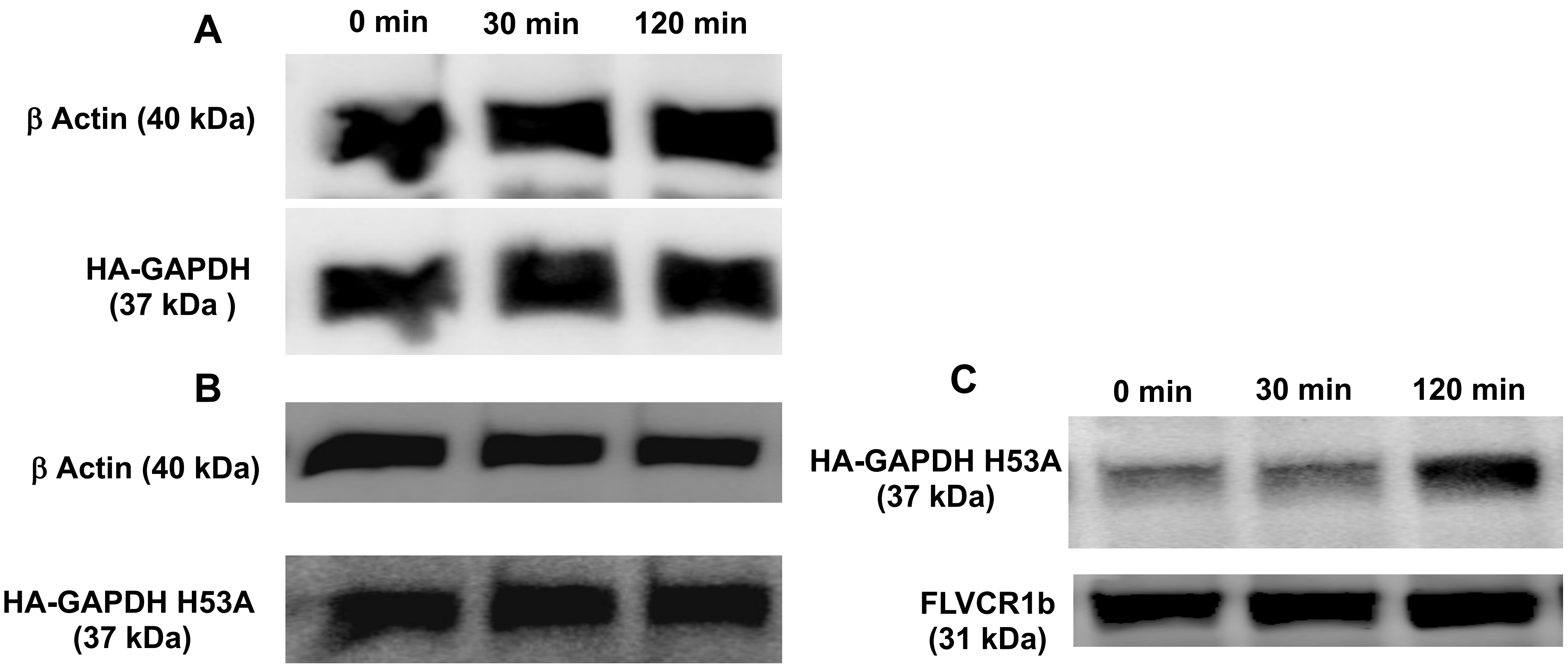


**Supplementary Figure 4. Time courses of FLVCR1b interaction with wild type or H53A HA-GAPDH in cells after stimulation of their mitochondrial heme biosynthesis. A**, **B** Reprentative Western blots showing the relative expression levels of β-actin, HA-GAPDH, or the HA-GAPDH H53A variant in supernatant samples (equal total protein) from transfected HEK293T cells after they had received δ-ALA/Fe and underwent further culture for the indicated times prior to harvest. **C** HEK293T cells expressing the HA-GAPDH H53A variant were harvested at the indicated time points after δ-ALA/Fe addition and cell supernatant samples (equal total protein) were subject to pulldown using anti-FLVCR1 antibody. Representative Western blot shows the relative levels of HA-GAPDH H53A and FLVCR1b proteins in the pulldown samples.

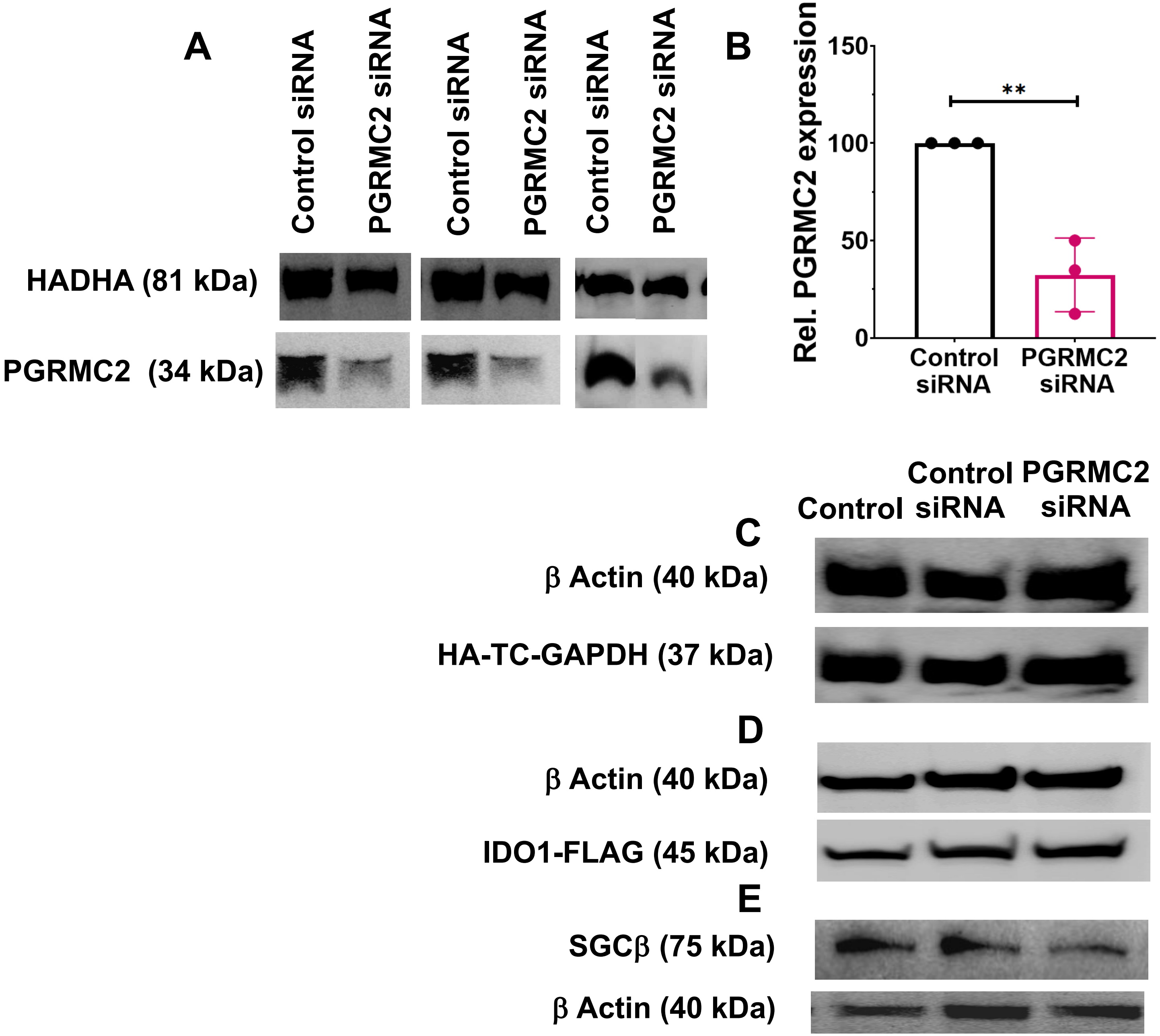


**Supplementary Figure 5. Impact of scrambled (control) or PGRMC2-targeted siRNA treatment on the expression levels of the indicated proteins in transfected HEK293T cells.** The images are from representative Western blots. Equal total protein amounts of each cell supernatant sample were run on SDS-PAGE and Western blotted and developed using antibodies against the indicated proteins or tags. **A** HADHA and PGRMC2 expression levels in response to the scrambled or targeted siRNA treatment. **B** Quantitation of the results from A based on band intensities. From three independent trials, mean +/- SD. Significance: ** $p < 0.01$ vs. the compared group based on a two tailed t-test. **C, D, E** Expression levels of the indicated proteins in cells that had been given vehicle, scrambled siRNA, or PGRMC2-targeted siRNA.

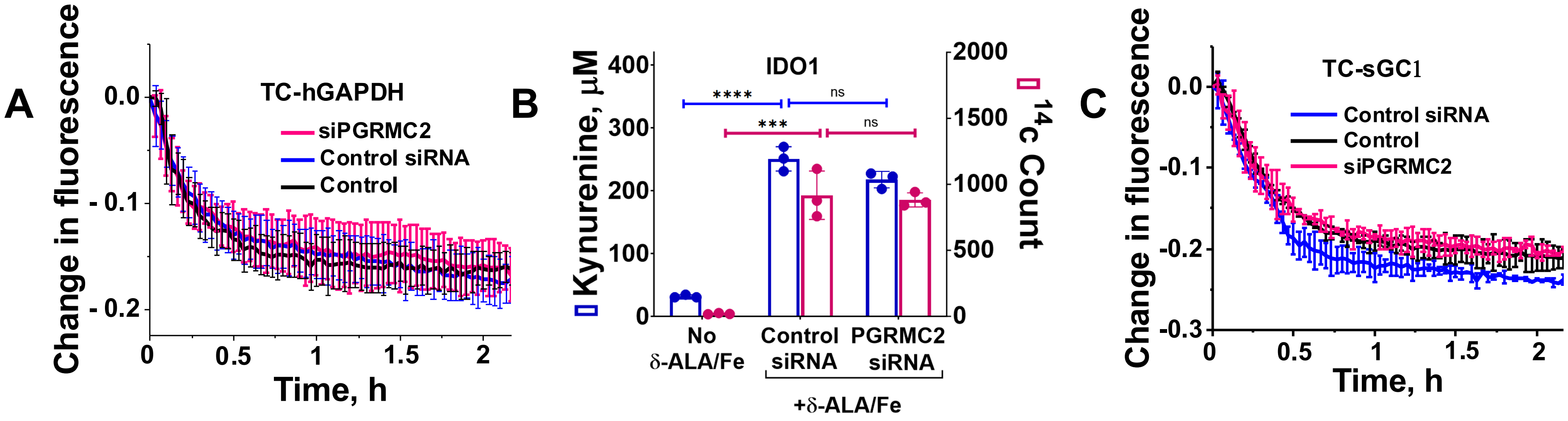


**Supplementary Figure 6. Knockdown of cell PGRMC2 protein expression does not impact the transfer of mitochondrial heme into TC-hGAPDH or the downstream GAPDH-dependent heme deliveries to IDO1 and sGCβ.** HEK293T cells underwent treatment with vehicle alone (control) or with scrambled (control siRNA) or PGRMC2-directed siRNA, and were transfected to express either TC-hGAPDH, IDO1, or TC-sGCβ. Cells were then given cold or $^{14}$C-radiolabeled δ-ALA/Fe and further cultured. **A** Time course of FlAsH-labeled TC-hGAPDH heme binding in live cells after they were given δ-ALA/Fe at Time = 0. **B** IDO1 activity in cells after δ-ALA/Fe addition as assessed by the Kynurenine product that accumulated in the cell culture fluid; and by the content of IDO1 $^{14}$C-heme, both as measured from IDO1 pulldown of cell supernatants (equal total protein) made from cells harvested 48 h after IDO1 transfection and addition of $^{14}$C-δ-ALA/Fe. **C** Time course of FlAsH-labeled TC-sGCβ heme binding in live cells after they were given δ-ALA/Fe at Time = 0. **A, C** Representative of three independent trials, data points are the mean +/- SD of triplicates. **B** Three independent trials, mean +/- SD. Significance: **** $p < 0.0001$, *** $p < 0.001$ vs. the compared group based on a one-way ANOVA. ns, not significant.

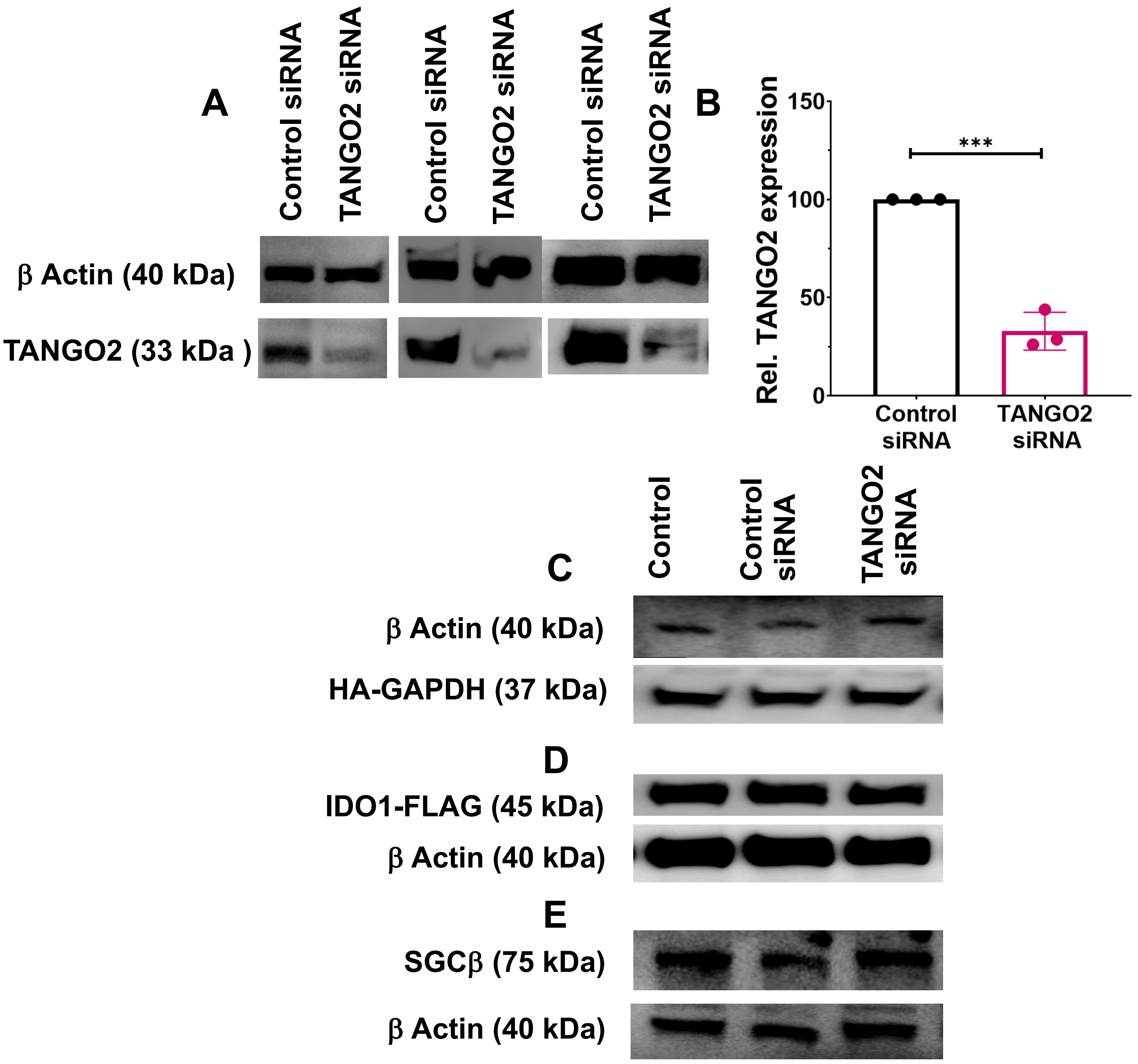


**Supplementary Figure 7. Impact of scrambled (control) or TANGO2-targeted siRNA treatment on the expression levels of the indicated proteins in transfected HEK293T cells.** The images are from representative Western blots. Equal total protein amounts of each cell supernatant sample were run on SDS-PAGE and Western blotted and developed using antibodies against the indicated proteins or tags. **A** β-actin and TANGO2 expression levels in response to the scrambled or targeted siRNA treatment. **B** Quantitation of the results from A based on band intensities. Three independent trials, mean +/- SD. Significance: *** $p < 0.001$ vs. the compared group based on a two-tailed t-test. **C, D, E** Expression levels of the indicated proteins in cells that had been given vehicle, scrambled siRNA (control), or TANGO2-targeted siRNA.

Table S2. Total soluble heme content in control and si-RNA treated HEK293T cell supernatants

Values are the mean +/- SD from 3 independent experiments

| Normal (pmol/mg protein) | siTANGO2 (pmol/mg protein) |
|---|---|
| 62 ± 2 | 66 ± 2 |

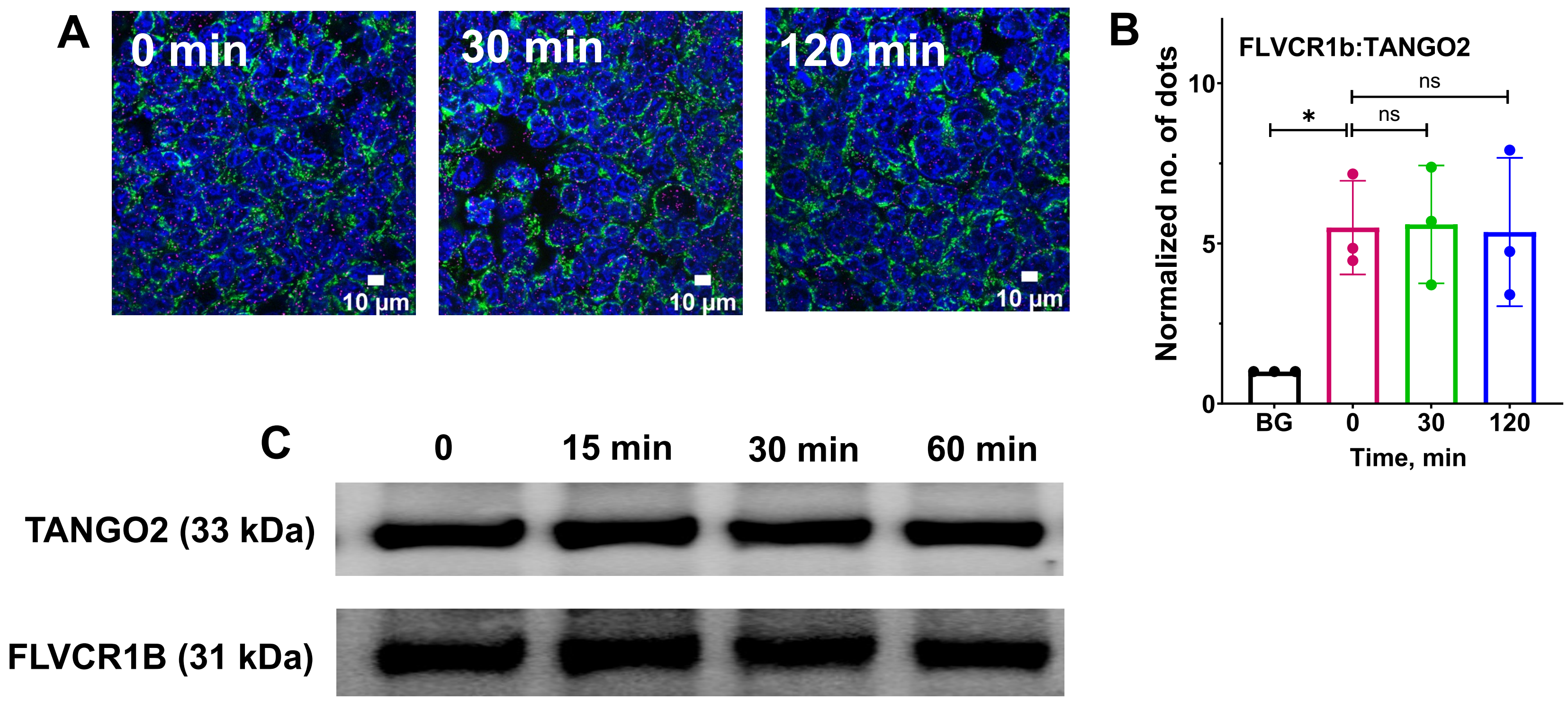


**Supplementary Figure 8. TANGO2 interaction with FLVCR1b in cells does not change after stimulation of their mitochondrial heme biosynthesis.** HEK293T cells were given δ-ALA/Fe to stimulate mitochondrial heme biosynthesis and then were harvested for processing at the indicated times. **A** PLA was used to assess TANGO2 interaction with FLVCR1b. Representative fluorescence microscope images of cells stained with DAPI against nuclei (blue), mitochondria (HADHA, green), and to show the TANGO2-FLVCR1b interaction by PLA (pink). **B** Quantification of the PLA results showing the extent of TANGO2-FLVCR1b interaction vs time after the δ-ALA/Fe addition. BG, the background PLA signal level. Three independent trials, mean +/- SD. Significance: * $p < 0.05$ vs. the compared group based on a one-way ANOVA. ns, not significant. **C** Representative Western blots showing the relative levels of FLVCR1b and TANGO2 proteins after pulldown with an anti-FLVCR1 antibody of the cell supernatant samples (equal total protein).

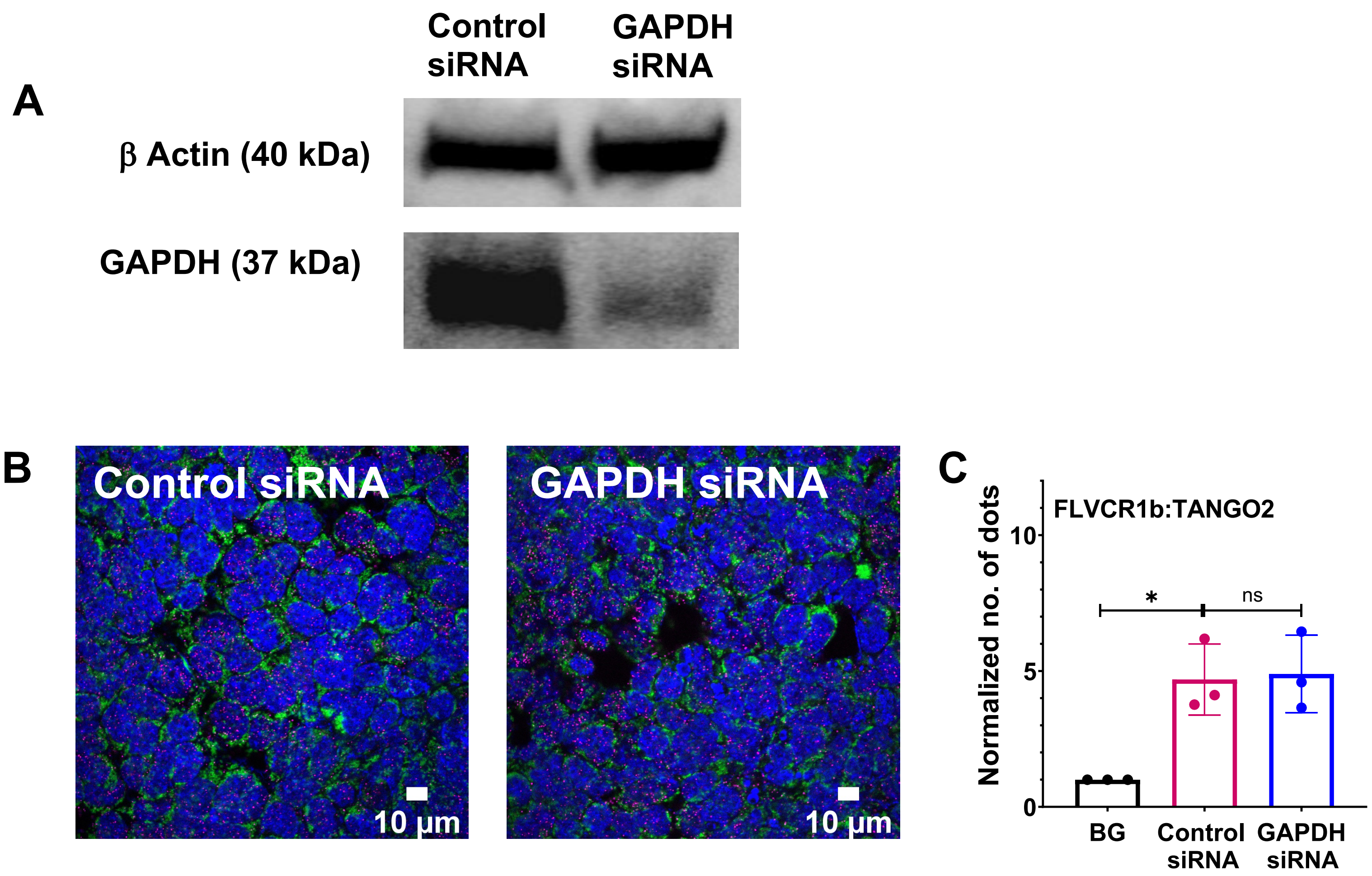


**Supplementary Figure 9. Impact of GAPDH knockdown on the TANGO2 interaction with FLVCR1b in cells.** HEK293T cells were treated with scrambled (control) or GAPDH-directed siRNA and then further cultured for 48 h before processing. **A** Representative Western blot showing knockdown of GAPDH expression in response to the siRNA treatment. **B** PLA was used to assess TANGO2 interaction with FLVCR1b. Representative fluorescence microscope images of cells stained with DAPI against nuclei (blue), mitochondria (HADHA, green), and to show the TANGO2-FLVCR1b interaction by PLA (pink). **C** Quantification of the PLA results showing the level of TANGO2-FLVCR1b interaction in the cells. BG, the background PLA signal level. Three independent trials, mean +/- SD. Significance: * $p < 0.05$ vs. the compared group based on a one-way ANOVA. ns, not significant.

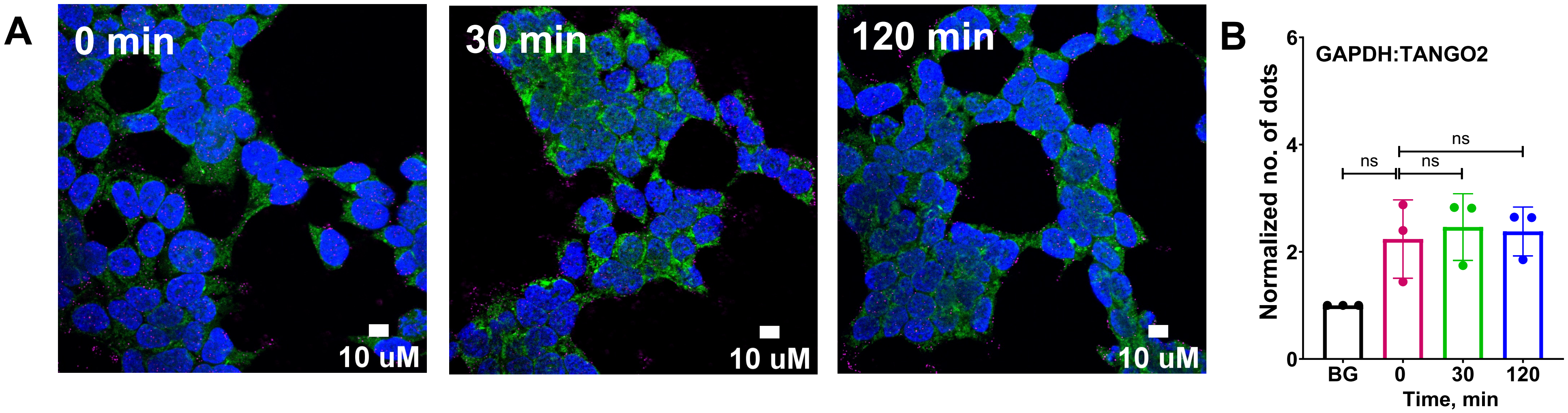


**Supplementary Figure 10. GAPDH interacts weakly with TANGO2 in cells.** HEK293T cells were given δ-ALA /Fe and incubated further for the indicated times before being processed. **A** PLA was used to assess the level of TANGO2 interaction with GAPDH. **A** Representative fluorescence microscope images of cells stained with DAPI against nuclei (blue), mitochondria (HADHA, green), and to show the TANGO2-GAPDH interaction by PLA (pink). **B** Quantification of the PLA results compares the level of GAPDH-TANGO2 interaction during the δ-ALA/Fe incubation. BG, the background PLA signal level. Three independent trials, mean +/- SD. Significance: one-way ANOVA. ns, not significant.

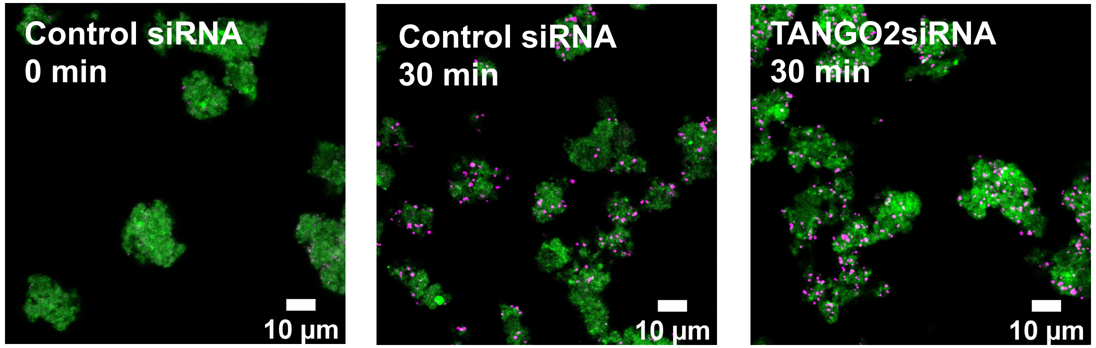


**Supplementary Figure 11. Cytosolic TANGO2 is not needed for the FLVCR1b-GAPDH interaction to increase in isolated mitochondria during stimulation of their heme biosynthesis.** Representative fluorescence microscope images of isolated mitochondria stained with antibodies against mitochondria (HADHA, green) and to show the level of GAPDH-FLVCR1b interaction by PLA (pink). The mitochondria were reisolated after having been incubated for the indicated times with δ-ALA plus mitochondria-free supernatants that were prepared either from control or TANGO2 knockdown HEK293T cells to stimulate heme biosynthesis.
